# *In Situ* Structure Determination of a Membrane Protein in Native Cellular Membranes by Proton-Detected Solid-State NMR

**DOI:** 10.1101/2025.10.28.685061

**Authors:** Huayong Xie, Weijing Zhao, Hang Xiao, Yan Zhang, Yang Shen, Yuefang Gan, Qiong Tong, Yongxiang Zhao, Huan Tan, Jun Yang

## Abstract

Determining the structure of membrane proteins within their native cellular membranes remains a substantial challenge in structural biology. In this study, we present a proton-detected solid-state NMR (ssNMR) approach, combined with an optimized random partial protonation (RAP) labeling strategy, to determine the high-resolution structure of the large-conductance mechanosensitive channel (MscL) directly within native *E.coli* membranes (backbone RMSD = 1.9 Å). Our approach effectively suppresses background protein signals and achieves high spectral resolution and sensitivity at moderate MAS frequencies (40–60 kHz) by differentially tuning amide and side-chain protonation levels. Using advanced recoupling schemes, we obtained chemical shift assignments of side-chain protons by 3D hCCH spectra and ^1^H-^1^H distance restraints from a series of 3D hNHH spectra. With 10% protonation in side-chains, the ^1^H signals displayed linewidths of approximately 50Hz, facilitating the extraction of 49 long-range distance restraints between amide and side-chain protons, which are crucial for structural convergence. Ambiguities in the assignment of weak signals corresponding to distance restraints were resolved by integrating 3D hNHH experimental data with CS-Rosetta structural modeling. The resulting structure reveals a well-defined pentameric assembly, with transmembrane helix packing consistent with that observed in detergent environments. This study demonstrates significant sensitivity advantages of ^1^H-detected over ^13^C-detected *in situ* ssNMR methods, highlighting the potential of ^1^H-detected ssNMR for the structure determination of a broad range of membrane proteins in native membranes.

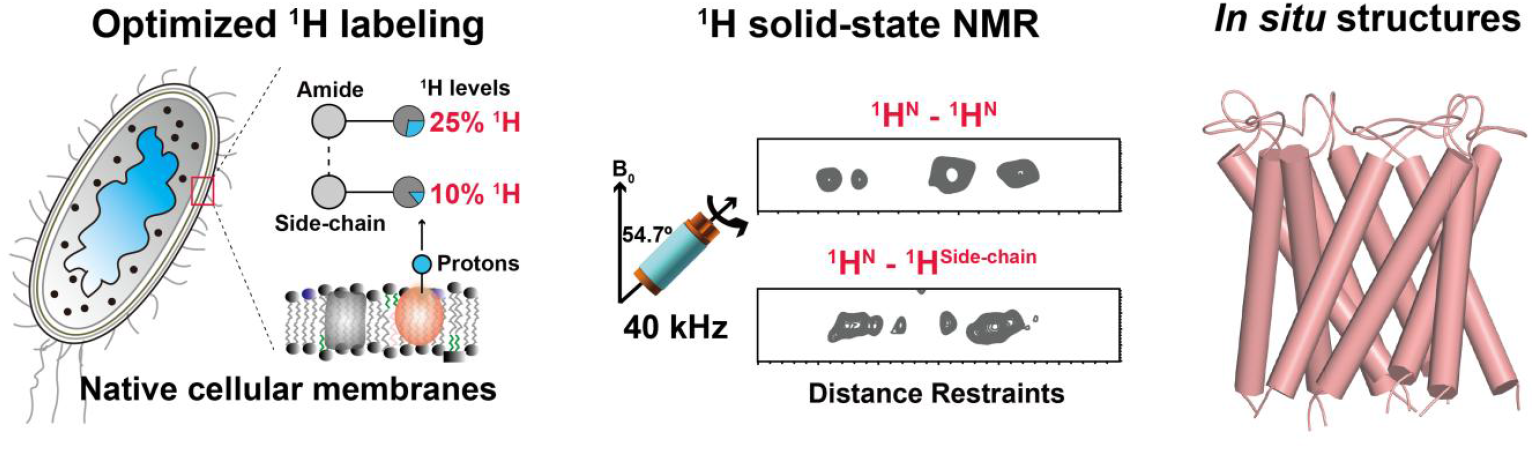

## INTRODUCTION

Membrane proteins play central roles in cellular processes, yet their precise three-dimensional structures, dynamic conformations, and functional activities are highly dependent on their native lipid environment^1,2^. The cellular membrane is not a homogeneous phospholipid barrier but rather a complex, dynamic, and regionally heterogeneous system composed of many lipid species, proteins and other components^3^. Although structural studies of membrane proteins in membrane-mimetic environments, such as detergent micelles or artificial liposomes, have provided valuable insights into their molecular mechanism^4–7^, these approaches carry an inherent risk of structural distortion^1,8,9^. Therefore, developing methods that enable direct, high-resolution structural determination of membrane proteins within native cellular membranes is essential for accurately elucidating their true molecular mechanisms^10–12^.

Substantial progress has been made in the *in situ* ssNMR analysis of membrane proteins within native environments^13–16^. Several strategies, including the use of deletion strains^17^, expression in “dual media”^18^, antibiotic treatment^19,20^ and isolation of membrane components^21^, have been developed to enhance the concentration of target proteins while suppressing background protein expression and labeling. These methods have enabled structural analysis of various membrane protein embedded in cellular membranes, such as LR11^22^, bR^23^, Mistic^24^, CsmA^25^, M2^26^, PagL^17,27^, T4SScc^28^, YadA^29^, ASR^21^, EGFR^30^, KcsA^31^, YidC^20^, BAM^32^ and LHCII^33^. Building these advances, our group has established a protocol for cellular membrane preparation that combines optimized expression conditions and bacterial strains, a dual media labeling approach, and selective removal of non-target membrane components^34^. This protocol achieved nearly complete suppression of background protein signals while maintaining high spectral sensitivity for the target protein, leading to the first high resolution 3D structure of AqpZ in inner membranes of *E. coli* determined using ^13^C-detected ssNMR spectra^10^.

However, the low sensitivity of ^13^C-detected ssNMR, resulting from lower gyromagnetic ratio of ^13^C compared to ^1^H, limits the widespread application of *in situ* ssNMR studies of membrane proteins^13,35^. To enhance ^13^C-detected spectral sensitivity, it is necessary to increase isotopically labeled protein samples to tens milligrams and to extend data acquisition times^36^. Moreover, this low sensitivity restricts the acquisition of spectra with dimensionality higher than 2D, which is an effective approach for resolving ambiguities in extraction of distance restraints by disentangling highly overlapped 2D ^13^C-^13^C spectra^37^.

^1^H-detected ssNMR has emerged as a powerful approach to overcome the limitations of ^13^C-based detection^38,35,36^. The high gyromagnetic ratio (γ) of protons inherently provides significantly higher sensitivity compared to ^13^C, theoretically by a factor of ∼8^36,39^. However, strong ^1^H-^1^H homonuclear dipole-dipole couplings cause substantial line broadening, historically limiting the use of ^1^H detection in solid-state biomolecular NMR due to compromised resolution and sensitivity^35,40^. Recent advances in fast magic angle spinning (MAS) probe technology and deuterated protein sample preparation have alleviated these dipole-dipole interactions, enabling the determination of more than ten biomolecular ssNMR structures based on ^1^H detection^41,35,42–44,41,45–48,37,49^. However, applying ^1^H-detected ssNMR for membrane protein structure determination in native cellular membrane remains highly challenging due to several factors^13^: (i) Background signal interference. Native membranes contain abundant background proteins and small molecules, whose ^1^H signals may overlap obscure those of the target protein. (ii) Limited spectral resolution and sensitivity. The inherent low abundance of membrane proteins in native membranes reduces spectral sensitivity. Additionally, the lack of effective deuterated protocol for membrane proteins in native membranes further limits ^1^H spectral resolution. (iii) Ambiguity in extraction of distance restraints. The narrow chemical shift dispersion of ^1^H (0–10 ppm) and the inability to definitively distinguish intra-from inter-molecular signals in native membranes introduce substantial ambiguities when extracting ^1^H-^1^H distance restraints, thereby reducing structural calculation accuracy.

To address these challenges, we established a comprehensive methodology for determining 3D structures of membrane proteins in native membranes by ^1^H-detected ssNMR spectra. This methodology includes a protocol for preparing cellular membrane samples, the corresponding 3D pulse sequences and an approach for solving ambiguities in extracting distance restraints. We employed the large-conductance mechanosensitive channel (MscL) as a model protein to demonstrate the feasibility of our methods^50,51^. MscL is a pentameric channel, with monomer containing two transmembrane helices in its X-ray crystal structure^52^. We have previously achieved nearly complete assignment of amide proton (^1^H^N^), ^13^C, and ^15^N chemical shifts for MscL in native *E. coli* inner membranes^53,54^. In this study, we modified our previous protocol in ^13^C detection for partial deuterated sample using a RAP^55–57^ (Reduced Adjoining Protonation) approach, which allowed us to obtain high resolution and high sensitivity ^1^H spectra for structure determinations. We established a 3D hCCH-CP-PC4 experiment for assigning chemical shifts of side-chain ^1^H and a 3D hNHH-RFDR/hNHH-MODIST scheme for extracting ^1^H-^1^H distance restraints. Using a series of 3D hNHH experimental data and CS-Rosetta structural modeling, we obtained 92 long-range ^1^H-^1^H distance restraints, including correlations between amide and side-chain ^1^H atoms. Based on these restraints, we calculated high resolution structures of MscL with a backbone RMSD of 1.9 Å.

## RESULTS

### Cellular Membrane Sample Preparations of MscL for High-quality ^1^H-Detected ssNMR Spectra

High spectral resolution and sensitivity are essential for protein structure determination by ^1^H-detected ssNMR spectra^35^. Under the fast MAS rate, both resolution and sensitivity can be optimized by adjusting H2O/D2O ratio in the expression media and fine-tuning protonation levels of proton back-exchange in RAP^55,56^. Additionally, elimination of background protein signals from ssNMR spectra is critical to accurately extract distance restraints. In our previous study, we established a cellular membrane sample preparation protocol that combines strain optimization, dual-media culture, and outer membrane separation to completely eliminate ^13^C and ^15^N labeled background proteins from the cellular membranes^10^. In this study, we examined whether this protocol could be adapted to prepare RAP-deuterated samples devoid of background protein signals for ^1^H-detected ^1^H-^15^N and ^1^H-^13^C correlated ssNMR spectra (**Figure 1a**). We prepared four membrane samples with varying deuteration levels in protein expression and in protonation back exchange (**Figure 1b-e** and **Table S1**). Additionally, to identify potential background protein signals, we reconstituted purified ^13^C,^15^N-labeled MscL with a 10% protonation level in POPC/POPG liposomes as a background-free reference.

**Figure 1.**
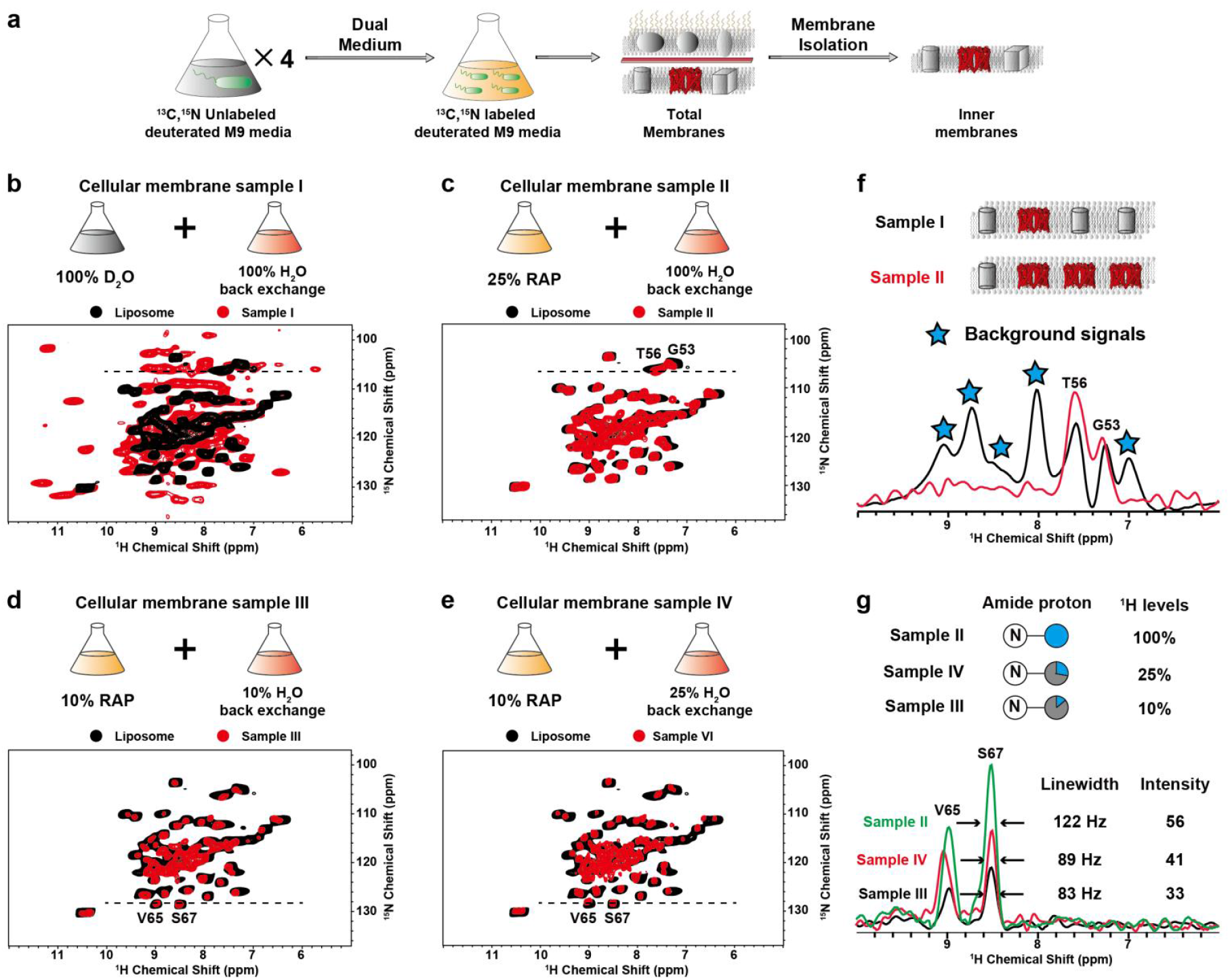
Screening of deuterium levels in cellular membrane sample preparations using the RAP technique to obtain high-quality ssNMR spectra. **(a)** Schematic workflow for preparing high-quality ^2^H, ^13^C, ^15^N-labeled MscL membrane samples, adapted from our previous study^34^. Cells were first cultured in deuterated M9 medium without ^13^C/^15^N labeling, then concentrated 4:1 into ^13^C/^15^N-labeled deuterated M9 medium for MaMscL expression induction, and finally inner membranes were collected via sucrose density gradient centrifugation. **(b-e)** The H_2_O/D_2_O ratio in the expression medium and the protonation level during proton back-exchange are critical for the resolution and sensitivity of ^1^H ssNMR spectra. To identify the optimal deuterization scheme, four cellular membrane samples (I - IV) with varying H_2_O/D_2_O ratios during expression and back-exchange were analyzed by 2D ^1^H - ^15^N HSQC spectra (red). Using reconstituted MscL in liposomes (10% protonation, black) as background-free reference, spectrum from Sample I shows substantial background protein signals while Samples II - IV are clean. **(f)** Comparison of 1D slices from the HSQC spectra (I, black; II, red) confirms effective background protein signals suppression in spectrum from Sample IV. Schematics above illustrate relative content of background proteins (gray) vs MscL (red). **(g)** 1D slices of the HSQC spectra from Samples II–IV were compared (II, green; III, black; IV, red). Sample IV exhibits a balanced signal-to-noise ratio and resolution. The amide protonation levels of the three samples are indicated above the 1D slices.

Sample I was prepared using our protocol (**Figure 1a**) following a conventional procedure to reduce ^1^H density of proteins: MscL were expressed in the medium containing U-[^2^H,^13^C]glucose and 100% D_2_O, followed by back-exchange in 100% H_2_O to protonate amide group and solvent-accessible sites. Its 2D ^15^N–^1^H HSQC spectrum is shown in **Figure 1b**. Unexpectedly, strong ^1^H signals of residues in all domains, including those in the hydrophobic transmembrane regions, were observed after proton back-exchange, possibly due to the large funnel-like solvent accessibility of its structure^52^. The spectrum showed additional peaks compared to that of the purified MscL reconstituted in proteoliposomes. After accounting for potential multi-bond polarization transfer and signal aliasing (**Figure S1A**), these extra peaks were assigned to background proteins. Their presences resulted from seriously impaired MscL expression in M9 medium containing 100% D_2_O, which increased the relative abundance of background proteins (**Figure S1B, C**).

To recover high level MscL expression, we prepared Sample II using a 75% deuterated RAP strategy: cells were grown in media containing U-[^2^H,^13^C]glucose and 25% H_2_O/75% D2O. To enhance ^1^H-^1^H correlation signals, partial deuterated protein was further back-exchanged in 100% H_2_O to fully protonate of amide ^1^H. As shown in **Figure 1c**, the partially diluted (100% ^1^H^N^ - 25% side-chain ^1^H (^1^H^SC^)) and U-[^13^C,^15^N]-MscL native membrane sample exhibited spectra with almost no background signals. A comparison of 1D slices from spectra of Samples I and II clearly demonstrates the elimination of background protein signals (**Figure 1f**). However, Sample II showed a ^1^H linewidth of ∼130 Hz in 2D ^1^H–^15^N HSQC spectra, indicating the need for further optimization of spectral resolution.

In our prior studies^53^, we acquired 2D ^1^H–^15^N correlation spectra of MscL at four deuteration levels (60%, 70%, 80%, and 90%) in RAP, observing average amide ^1^H linewidths of 146, 135, 126, and 105 Hz, respectively. Based on these results, a 10% protonation level was selected for ^1^H dilution (Sample III, **Figure 1d**). Although Sample III demonstrated a ∼30 Hz reduction in linewidth compared to Sample II, its amide signal intensity was only ∼50% of that in Sample II. Considering the trade-off between sensitivity and resolution, Sample II was chosen for acquiring 3D hNHH spectra to obtain amide-amide (^1^H^N^–^1^H^N^) distance restraints.

To improve side-chain ^1^H resolution and obtain more side-chaindistance restraints (^1^H^N^-^1^H^SC^), we designed Sample IV (**Figure 1e**) using a 10% protonation in RAP followed by back-exchange in 25% H_2_O. This resulted in 25% amide protonation at water-accessible sites and 10% side-chain protonation (**Figure 2a**). Compared to Sample II, Sample IV showed a ∼30% decrease in amide signal-to-noise ratio but a ∼20% improvement in its resolution (**Figure 1g**), offering only limited advantage for extracting amide-amide (^1^H^N^-^1^H^N^) distance restraints in 3D hNHH experiments.

**Figure 2.**
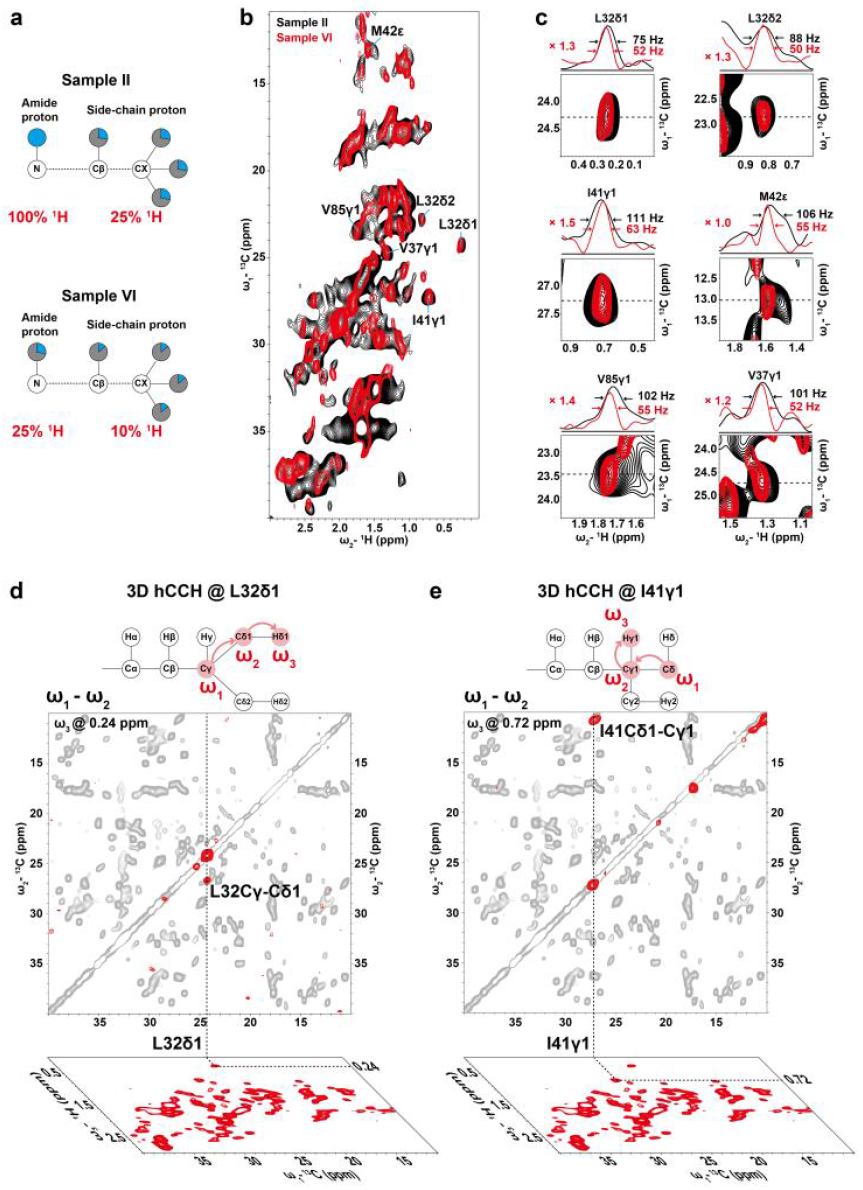
Screening optimal deuterium levels for side-chain ^1^H resolution to facilitate resonance assignment and distance restraint extraction. **(a)** Amide and side-chain protonation levels in Samples II and IV. **(b)** Overlay of 2D hCH spectra from Sample II (black) and Sample IV (red). Spectral regions and 1D slices for six representative residues are expanded in panel **(c)**, showing that Sample IV exhibits a ∼50 Hz reduction in side-chain ^1^H linewidth compared to Sample II, albeit with an approximately 20% decrease in signal-to-noise. Sample IV was therefore selected for subsequent resonance assignment and distance restraint extraction. **(d-e)** Strategy for assigning side-chain proton chemical shifts using 3D hCCH spectra. Since nearly all cross-peaks in the 2D ^13^C–^13^C spectrum of reconstituted MscL in proteoliposomes have been previously assigned^54^, each w_1_(^13^C)–w_2_(^13^C) plane of the 3D hCCH spectrum can be overlaid with the 2D ^13^C–^13^C reference (black) to assign the w_1_(^13^C) and w_2_(^13^C) chemical shifts of a target peak. The w_3_(^1^H) dimension corresponds to the proton directly bonded to the carbon atom of w_2_(^13^C), allowing proton chemical shift assignment once the w_2_(^13^C) atom is identified. For example, the 3D hCCH cross-peak at (24.2, 26.6, 0.24) ppm shows a w_1_–w_2_ projection overlapping with the L32 C_γ_–C_δ1_ correlation in the 2D ^13^C–^13^C spectrum, enabling assignment of the 0.24 ppm signal to L32 Hδ1. Thus, the peak is assigned to L32 Cγ–Cδ1–Hδ1 **(d)**. Similarly, the peak at (10.5, 27.0, 0.72) ppm is assigned to I41 Cδ1–Cγ1–Hγ1 **(e)**.

Surprisingly, Sample IV exhibited a remarkable improvement in side-chain ^1^H resolution. A comparison of 2D hCH spectra between Samples II and IV revealed significantly reduced side-chain ^1^H linewidths in Sample IV (**Figure 2b**). Quantitative analysis showed side-chain ^1^H linewidths of ∼101 Hz in Sample II and ∼54 Hz in Sample IV, despite a ∼20% decrease in signal-to-noise ratio (**Figure 2c**). Therefore, when considering both sensitivity and resolution, Sample IV offers distinct advantages for acquiring side-chain distance restraints.

### Side-Chain ^1^H Chemical Shift Assignments in MscL Membrane Samples

Site-specific resonance assignment is essential for structural determination by NMR. In a previous study, we assigned ^13^C and ^15^N chemical shifts for 73 out of 101 residues of MscL in cellular membranes using ^13^C-detected spectra^54^, and 69 ^1^H^N^ using ^1^H detected spectra^53^. In this study, we further assigned side-chain ^1^H of MscL in cellular membranes to extract side-chain distance restraints. Our general strategy involves correlating detected ^1^H spins with nearby heteronuclei (typically ^13^C and ^15^N). Using this approach, we assigned side-chain ^1^H chemical shifts of MscL using 3D hCCH spectra, which encodes aliphatic side-chain ^1^H and ^13^C spins for directly bonded pairs (e.g., Hβ–Cβ, Hγ–Cγ; schematics in **Figure 2d, e**). In the 3D hCCH experiments, we employed the CP sequence^58^ for the initial ^1^H-^13^C transfer, the PC4 method^59^ for ^13^C-^13^C transfer and the INEPT sequence^60^ for the final ^13^C-^1^H transfer (**Figure S2A**). The PC4 method enhances ^13^Cα-^13^Cβ correlations, achieving a 50% improvement over commonly used DARR and TOBSY recoupling^59^. The ^1^H-^13^C INEPT sequence specifically transfer polarization of ^13^C to directly bonded protons, enabling the assignment of ^1^H chemical shifts attached to ^13^C. The 1D hCCH-CP-PC4 spectrum exhibits an 8-fold higher signal-to-noise ratio compared to the 1D hCCH-INEPT-TOBSY spectrum (**Figure S2B, C**).

Sample preparations, including both fully protonated and proton-diluted variants, exhibited high reproducibility, as evidenced by consistent ^13^C and ^15^N chemical shifts across samples^53^. Thus, we were able to assign peaks in the 2D ^13^C-^13^C plane (**Figure 2d, e**, red) from 3D hCCH by superposing them on well-resolved and assigned ^13^C-^13^C correlations from proteoliposome samples (black)^54^. The chemical shifts of ^1^H dimension can be assigned using the assigned ^13^C-^13^C correlations. For example: (i) The hCCH peak at (24.2, 26.6, 0.24) ppm, projected at (24.2, 26.6) ppm, overlapped with the L32 Cγ-Cδ1 cross-peak, thereby assigning L32 Hδ1 at 0.24 ppm. (ii) The peak at (10.5, 27.0, 0.72) ppm coincided with I41 Cδ1-Cγ1, assigning I41 Hγ1 at 0.72 ppm. A total of 53 side-chain ^1^H resonances were assigned using this 3D hCCH spectra (**Figure S3 and Table S2**).

### ^1^H^N^ - ^1^H^N^ and ^1^H^N^ - ^1^H^SC^ Distance Constraints extracted from ssNMR spectra

To determine the 3D structure of proteins by ssNMR, a sufficient number of distance restraints are required for structure calculations^61^. In this study, we acquired five 3D hNHH spectra of different samples using various recoupling schemes to enhance correlation signals corresponding to amide–amide and amide–side-chain proton proximity constraints (**Table S1**).

First, to enhance amide–amide correlations, we acquired 3D hNHH spectra of Sample II using radio frequency-driven recoupling (RFDR)^62^ scheme with an optimized mixing time of 2.0 ms and 4.0 ms for ^1^H-^1^H recoupling (**Figure S4A**). Theoretical simulations indicate that 2–3 ms is the optimal time window for RFDR recoupling (**Figure S5, a-c**). When the mixing time exceeds 3 ms, a significant attenuation in the intensity of long-range backbone ^1^H–^1^H correlation signals is observed. This attenuation is attributed to polarization leakage from backbone to side-chain protons, which is a consequence of the broadband nature of RFDR combined with the presence of aliphatic protons in the [25% ^1^H,^13^C,^15^N]-MscL cellular membrane sample. Consequently, a 2.0 ms RFDR mixing time was selected for 3D hNHH spectra focusing on backbone Second, to enhance the spectral intensity of the signals corresponding to backbone ^1^H–^1^H long-range correlations in Sample II, we implemented the band-selective MODIST^63,64^(Modest Offset Difference Internuclear Selective Transfer) recoupling sequence, which restricts polarization leakage to side-chain protons (**Figure S4B**). As illustrated in **Figure S4C**, MODIST provides high frequency selectivity, effectively suppressing dipolar truncation and relay effects. This selectivity also simplifies NMR spectra by excluding undesired correlations, thereby improving the reliability of assignments of the signals corresponding to long-range distance assignments (**Figure 3a**, green). 1D ^1^H traces extracted from 3D hNHH spectra acquired with MODIST (**Figure S4D**) confirmed reduced polarization transfer to side chains, thereby preserving backbone magnetization and strengthening backbone ^1^H–^1^H cross-peaks. Notably, several long-range correlations that were weak or absent in hNHH spectra using RFDR recoupling appeared under the MODIST schemes, increasing the number of usable distance restraints (**Figure 4a, gree**n). Theoretical simulations (**Figure S5, d-f**) signals (**Figure 3a**, blue), and a 4.0 ms mixing time was used for spectra focusing on side-chain signals (**Figure 3b**, blue). indicated that MODIST requires mixing times longer than 4 ms for optimal efficiency; we therefore used 6.0 ms and 12.0 ms for 3D hNHH data acquisition. Subsequent spectral analysis identified 27 additional long-range restraints beyond those obtained with RFDR (**Figure 4a**, green).

**Figure 3.**
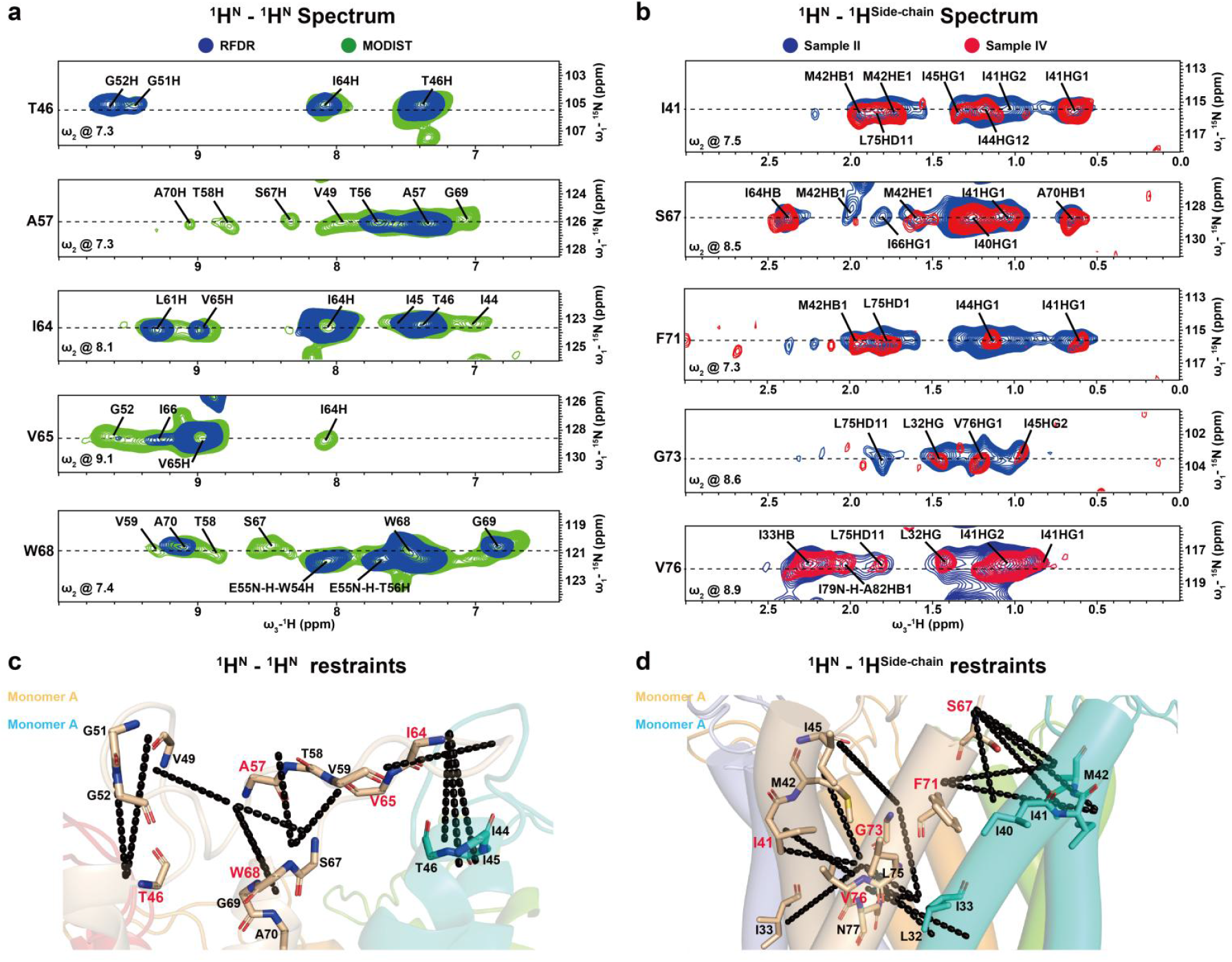
Extraction of distance constraints from ssNMR spectra using different recoupling schemes and sample preparation methods. **(a)** Overlaid 3D hNHH spectra of Sample II acquired with RFDR (2.0 ms, blue) and MODIST (6.0 ms, red) recoupling schemes, annotated with assigned peaks. **(b)** Comparison of 3D hNHH spectra from Sample II (RFDR 4.0 ms, blue) and Sample IV (MODIST 4.0 ms, red), with assigned peaks labeled. **(c)** ^1^H^N^-^1^H^N^ distance constraints mapped onto the lowest-energy CS-Rosetta structure, highlighting loop-helix interfaces. **(d)** ^1^H^N^-^1^H^SC^ distance constraints mapped onto the same structure, concentrated at helix-helix packing interfaces.

**Figure 4.**
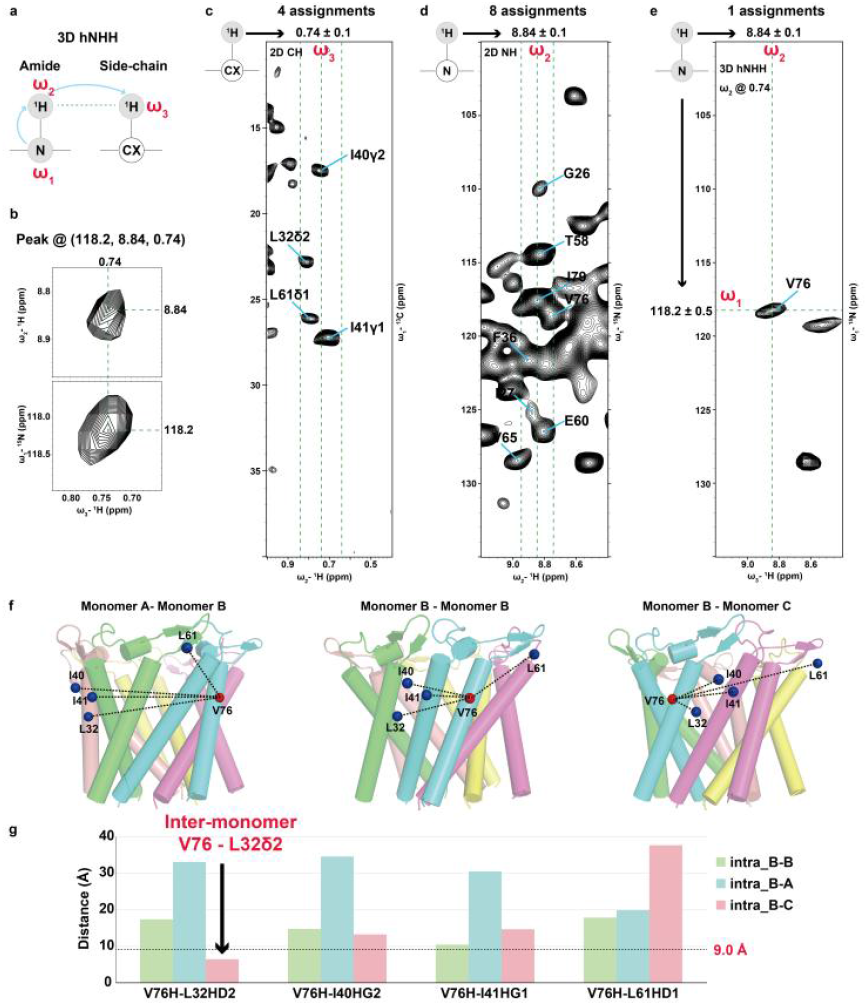
Resolving ambiguity in assignments of spectral signals corresponding to distance constraints by using 3D spectra and CS-Rosetta modeling. **(a)** Magnetization transfer pathway in the 3D hNHH experiment used for obtaining distance constraints. **(b)** w_2_-w_3_ and w_1_-w_3_ planes of the 3D hNHH peak at (118.2, 8.84, 0.74) ppm. **(c)** Four assignment candidates for w_3_@0.74 ppm (±0.1 ppm tolerance) mapped onto a 2D hCH spectrum. **(d)** Eight candidates for w_2_@8.84 ppm (±0.1 ppm tolerance) in a 2D hNH spectrum. **(e)** Combined filtering using w_1_ (118.2 ppm) and w_2_@8.84 ppm dimensions uniquely identifies V76. **(f)** Distance validation for four assignment candidates at monomer B-monomer A interface, monomer B interior, and monomer B-monomer C interface within the CS-Rosetta-derived pentameric structure. **(g)** Distance distribution analysis: Only inter-monomer V76 H^N^ - L32Hδ2 satisfies the 9.0-Å cutoff.

Third, to enhance amide–side-chain correlations, we acquired 3D hNHH spectra of Sample IV using RFDR scheme for ^1^H-^1^H recoupling at 4.0 ms mixing time (red). Compared to Sample II under the same conditions, Sample IV exhibited markedly improved side-chain resolution, which reduced assignment ambiguities for side-chain protons. By integrating assignments from both Samples II and IV, we obtained 49 long-range ^1^H^N^-^1^H^SC^ distance restraints (**Figure 4b**, red).

In the assignments of distance restraints, ^1^H chemical shifts were used to identify amino acids. To accurately assign the ^1^H signals from MscL, it was necessary to exclude the ^1^H signals originating from lipids and background proteins. We assigned ^1^H chemical shifts of lipids in *E.coli* membranes as they appear in dipolar-based spectra, and excluded signals with chemical shifts similar to these lipids in the assignments of signals for distance restraints of proteins (**Figure S6** and **Table S3**). Background protein ^1^H signals were also excluded, since these proteins are randomly distributed around the target protein and, many are located too far from MscL to generate detectable correlations with MscL^3,10,11,39,65^. Furthermore, the similarity of chemical shifts among atoms of different amino acids introduces multiple possible assignments for distance restraint peaks, introducing ambiguity that diminished the quality of structural calculations.

To address this challenge, we employed two complementary strategies. The first strategy involved the use of 3D spectra. Thanks to the high sensitivity of ^1^H detection, we were able to acquire 3D hNHH spectra for assigning distance restraints (∼71 hours acquisition time) (**Figure 4a and Table S1**). As an example, the peak at (118.2, 8.84, 0.74) ppm in the 3D hNHH spectrum illustrates how 3D data significantly reduces ambiguity in distance restraint peak assignments(**Figure 4b**). For the w3 @ 0.74 ppm dimension, the 2D hCH spectrum shows four possible assignments: I40γ2, L32δ2,L61δ1 and I41γ1 (**Figure 4c**). Meanwhile, in the w2 @ 8.84 ppm dimension, the 2D hNH spectrum shows eight possibilities: G26, T58, I79, V76, F36, I27, E60, and V65 (**Figure 4d**). Therefore, the peak at (8.84, 0.74) ppm has 32 (4 × 18) possible assignments. However, in the 3D hNHH spectrum, the ((w1, w2)) dimensions correspond to directly bonded N-H amide pairs. Using the w1 @ 118.2 ppm dimension to filter the w2 @ 8.84 ppm possibilities reduces the ambiguity from eight to one, with only V76 satisfying the chemical shift criteria (**Figure 4e**). Consequently, the ambiguity in the assigning the distance restraint peak at (8.84, 0.74) ppm decreases from 32 to 4, greatly improving the confidence of the assignment.

In oligomeric complexes, intra-subunit and inter-subunit are indistinguishable based solely on chemical shifts. For example, in the pentameric MscL, the four remaining assignments for the peak at (8.84, 0.74) ppm must be considered in three contexts: intra-subunit, clockwise inter-subunit (B–A), and counterclockwise inter-subunit (A–B), resulting in a total of (4 × 3 = 12) possible assignments. In *in vitro* studies using artificial liposome samples, this ambiguity can be resolved by using differential labeling strategies[ref], such as labeling half of the subunits with ^13^C and the other half with ^15^N/^19^F. in this case, ^13^C-^15^N or ^13^C-^19^F correlation peaks arise exclusively from inter-subunit contacts, effectively distinguishing intra- and inter-subunit interactins^66–68^. However, this approach is not applicable to in *in situ* study, where differential labeling cannot be implemented. To resolve this ambiguity, we used a monomeric structural model produced by CS-Rosetta. We quantitatively evaluated the four possible assignments within the pentameric MscL model, examing them at the monomer B–monomer A interface, within monomer B, and at the monomer B–monomer C interface (**Figure 4f**). Using a 9.0 Å distance cutoff^48,69^, only the inter-subunit (monomer B–monomer C interface) V76H^N^ - L32Hδ2 pair met the spatial proximity criterion (**Figure 4g**).

By integrating these two strategies, we successfully assigned distance restraints between amide protons and side-chain protons. The same approach is also applicable for assigning distance restraints between amide protons (**Figure S7**)

### Structural Calculations and Comparative Analysis

By using the five 3D hNHH spectra described above, we obtained a total of 43 long-range ^1^H^N^-^1^H^N^ distance restraints, 49 long-range ^1^H^N^-^1^H^SC^ distance restraints (**Figures S8, S9 and Table S4**), and 186 inter-residue restraints (**Table S5**). These restraints, together with 134 backbone angle restraints predicted by TALOS+, were incorporated into Xplor-NIH for structure calculation, yielding 1000 MscL structures. The pentameric state of MscL was validated by cross-linking experiments (**Figure S10**). The backbone RMSD of the top 10 lowest-energy pentameric structures was (1.9 ± 0.4) Å (**Table S6**), indicating good structural convergence.

To evaluate the contribution of side-chain proton restraints, we repeated the calculations after removing the 48 long-range ^1^H^N^-^1^H^SC^ distance restraints. Following the same protocol, the backbone RMSD of the top 10 lowest-energy pentameric structures increased to 3.8 ± 0.6 Å (**Figure 5a**), demonstrating that side-chain restraints play a critical role in enhancing structural accuracy. But how do these side-chain proton distance restraints exert such an effect? Mapping the distance restraints onto the structure reveals that while ^1^H^N^-^1^H^N^ restraints leave certain gaps in the interhelical regions, the long-range ^1^H^N^-^1^H^SC^ restraints are predominantly distributed between helices, effectively filling these gaps (**Figure 5b and Figure S9**). This clearly illustrates the crucial role of side-chain distance restraints in constraining interhelical distances during structure determination.

**Figure 5.**
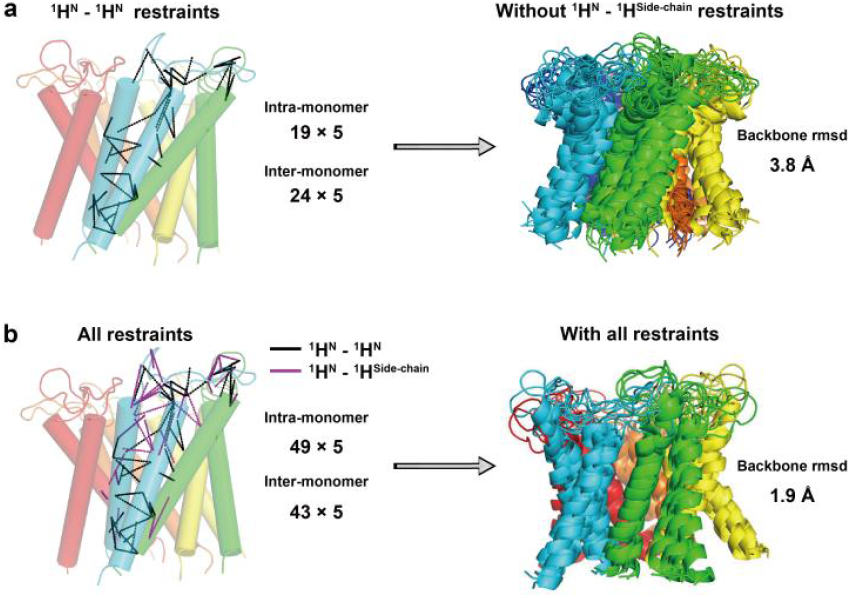
Critical role of ^1^H^N^ - ^1^H^SC^ distance constraints in defining MscL structural convergence. **(a)** Structural ensemble (10 lowest-energy models) calculated using only 43 long-range ^1^H^N^ – ^1^H^N^ restraints, showing a backbone RMSD of 3.8 Å. The spatial distribution of these ^1^H^N^–^1^H^N^ restraints is mapped onto the lowest-energy ssNMR structure. **(b)** Structural ensemble (10 lowest-energy models) obtained using both ^1^H^N^–^1^H^N^ and ^1^H^N^– ^1^H^SC^ restraints (92 in total), yielding a significantly improved backbone RMSD of 1.9 Å. The full set of long-range restraints is shown mapped onto the lowest-energy structure, illustrating the enhanced convergence enabled by including ^1^HN - ^1^H^SC^ restraints.

Since a pentameric structure of MscL was previously resolved in detergent micelles (PDB ID: 4Y7K) by X-ray crystallography^52^, we compared it with our structure determined in native membranes to assess environmental influences on MscL conformation (**Figure 6**). At the secondary structure level, the transmembrane helix composition remains consistent between environments–both structures feature two long transmembrane helices connected by an extended loop (**Figure S11A**). The tertiary structures are also highly similar: the arrangement of the two transmembrane helices is largely conserved, as reflected in a backbone RMSD of 3.3 Å between the two structures (**Figure 6a**). This suggests that the variations in secondary structure do not significantly alter the overall packing, indicating that tight hydrophobic packing within the MscL helix bundle confers considerable resistance to environmental perturbations. Quaternary structural analysis confirms that both structures form pentamers, with similar helix-helix interfaces composed of TM1 and TM2 in each case.

**Figure 6.**
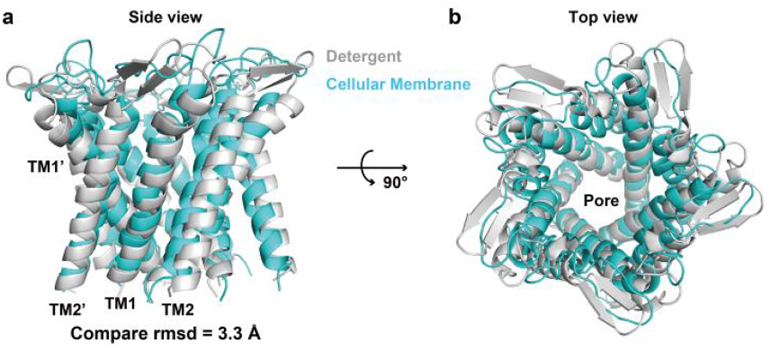
Comparison of the MscL structures determined by ssNMR in native membranes and by X-ray crystallography in detergent. **(a)** Side view of the ssNMR (cyan) and X-ray (gray) structures reveals highly consistent transmembrane helix tilts between the two environments, with only subtle variations. **(b)** Top view illustrating a well-conserved central pore architecture, demonstrating nearly identical pore in both structural models.

We further examined the tilt angles of the two transmembrane helices across both environments (**Figure S11B**). Owing to the limited number of distance restraints, the tilt angles derived for the native membrane structure exhibited some uncertainty. Statistical analysis of the top 10 lowest-energy structures was performed, and the mean tilt angles with standard deviations (σ) were calculated(**Figure S11B**). Using a 2σ threshold (99% confidence interval), TM2 helices showed tilt angles outside this range. While the detergent environment subtly perturbs the tilt angles of MscL transmembrane helices, the core structure remains largely stable (**Figure 6b**). This minimal structural alteration does not impair protein function, supporting our earlier findings that MscL activity is primarily regulated through dynamic mechanisms rather than large conformational changes^53^.

## DISCUSSION

This study demonstrates the remarkable advantages of *in situ* ssNMR based on ^1^H detection over ^13^C detection, exhibiting the great potential of ^1^H-detected *in situ* ssNMR in structure determination of membrane proteins in native membrane. The high gyromagnetic ratio (γ) of protons confers intrinsically higher sensitivity compared to ^13^C (theoretically by a factor of ∼8)^36,39^. This enhancement translates into two major practical benefits: drastically reduced sample requirements and significantly shortened acquisition times. Specifically, the native membrane sample amount decreased from ∼50 mg in previous ^13^C detection study^10,34,34,70^ to ∼10 mg for ^1^H detection in this study. A 2D ^13^C-^13^C correlation experiment for extracting ^13^C-^13^C distance restraints of AqpZ required ∼2 weeks^10,11^, whereas a 3D NHH experiment of MscL to extracting ^1^H-^1^H distance restraints in this study took only ∼3 days (**Table S1**). Additionally, the higher dimensional spectra afforded by ^1^H detection significantly enhances spectral resolution, enabling more unambiguous resonance assignments. The shortened acquisition times not only improves experimental efficiency but also reduces the risk of sample degradation during extended measurements, thereby facilitating studies of dynamic processes or less stable membrane proteins.

For membrane protein structure determination by *in situ* ssNMR in native membrane, the primary and most critical requirement is the suppression of background signals^13,34,8^. Since distance restraints for structure calculation generally derive from weak NMR signals, complete suppression of background protein signals in ssNMR spectra is essential for accurate extraction of these restraints. Two main strategies have been proposed to suppress background protein signals in cellular membrane sample preparations. The first strategy involves increasing the relative content of target proteins within cellular membranes^14,34^. Approaches aligned with this strategy include optimizing target protein expression^21,34^, removing non-target cellular membrane components^17,21,25^ and enriching target proteins by His-tag^16,21^. The second focuses selectively suppressing isotopic labeling of background proteins. To this end, methods such as optimizing expression strains, using gen-deleted strains^17^ and dual expression media^21^, and antibiotic treatment^19^ have been tested. In our recent studies, we established a protocol for preparation of cellular membrane samples by combining several elements of these two strategies^34^. Using this protocol, we obtained high-quality ^13^C and ^15^N ssNMR spectra of membrane proteins with minima interference from background protein signals and determined 3D structures of AqpZ^11^ and *Bj*SemiSWEET^10^ in *E. coli* inner membranes.

Suppression of background signals in ^1^H detection differs fundamentally from that in ^13^C detection. Unlike ^13^C detection, where background protein signals can be suppressed by isotopic selectively unlabeling, the ^1^H signals from target and background proteins cannot be directly distinguished by isotopic labeling during sample preparations. In this study, we filtered ^1^H species of target proteins by transferring magnetization from ^13^C and ^15^N nuclei of the labeled target proteins to their attached ^1^H nuclei in 3D hCCH and hNHH experiments (**Figure S2 and S4**). The effectiveness of these experiments depends critically on the complete suppression of ^13^C and ^15^N labeling of background proteins in cellular membranes. When only target proteins are ^13^C and ^15^N labeled in cellular membranes, the ^1^H signals of the target proteins can be cleanly filtered in 3D hCCH and hNHH. Therefore, with modifications in accordance with deuterated expression media in RAP method, the cellular membrane sample preparation protocol developed for ^13^C detection can be adapted for ^1^H detection. In this modified RAP-based protocol, utilizing expression media containing either 10% H_2_O/90% D_2_O or 25% H_2_O/75% D_2_O, the ^13^C and ^15^N labeling of background proteins is effectively suppressed. Consequently, their ^1^H signals can be indirectly suppressed due to the absence of corresponding ^13^C or ^15^N labels.

Another essential requirement for structure determination by ssNMR is that spectral resolution and sensitivity must be sufficiently high to enable accurate resonance assignments and extraction of distance restraints. A critical factor influencing the quality of ^1^H detected ssNMR spectra is the suppression of strong ^1^H-^1^H dipolar coupling present in solid proteins^35^. Two effective approaches to achieve high-resolution ^1^H spectra are extensive sample deuteration and high-speed MAS rates ranging from 40 to 160 kHz^35,71–73^. Although ssNMR structure determination using fully protonated samples exhibits significant advantages, such as requiring sub-milligram sample quality and providing abundant side-chain distance restraints^37,42^, it remains challenging for membrane protein structure determinations due to poor ^1^H chemical shift dispersion of membrane proteins. For example, ^1^H chemical shift dispersions for α-helical membrane proteins are only 3-4 ppm for amide protons (**Figure 1**) and 1-2 ppm for methyl protons (**Figure 2**), while ^1^H linewidths of amide and methyl protons are ∼ 180 and 150 Hz, respectively, at 100 kHz MAS rate in fully protonated protein samples^57,35,37^. This limited dispersion results in substantial signal overlap, especially in the methyl proton region, making resonance assignments difficult. The approach presented in this work combines ^1^H-detected ssNMR at fast MAS of 40 kHz with a RAP sample preparation strategy, effectively overcoming the limitations of H/D back-exchange in the transmembrane domains of membrane proteins. This enables structural studies to be extended to a broader range of membrane proteins. In contrast, H/D back-exchange of transmembrane domains of fully deuterated proteins have to use an unfold-refold procedure^57,46,47^, this limits its application to the membrane proteins without an available unfold-refold protocol. The key power of RAP lies in its fine control over deuteration levels, allowing an optimal trade-off between resolution and sensitivity tailored to specific experimental goals, including chemical shift assignments and distance restraints extraction^55–57^. This versatility makes RAP highly suitable for various phase of structural study. By enabling tunable protonation levels in both backbone and side-chain ^1^H species, RAP facilitates the acquisition of critical side-chain constraints even at moderate MAS frequencies. For instance, by using protonation levels below 10%, such as 5%, RAP achieves methyl ^1^H linewidths narrower than 50 Hz^55,56^, significantly improving signal resolution in the methyl regions and enabling structure determination of larger membrane proteins than MscL.

Currently, three membrane protein structures, including AqpZ^11^, *Bj*SemiSWEET^10^, and MscL, have been determined within native cellular membranes using *in situ* ssNMR. These structures exhibit varying degrees of sensitivity to their membrane environments. Membrane proteins with tightly packed transmembrane domains, such as AqpZ, ASR^21^ and MscL, are less affected by membrane composition. Their structures determined in native membranes closely resemble those resolved in detergent micelles or synthetic liposomes. In contrast, for membrane proteins with loosely folded transmembrane domains, like *Bj*SemiSWEET sugar transporter homodimer, show the conformational state and dynamics that are highly sensitive to the membrane environments. Functional conformational states of these proteins can only be captured within native membranes, which is essential for understanding fundamental biological mechanisms such as gating and transport.

The comprehensive methodology present in this study, which integrates a protocol for preparing partially deuterated membrane protein samples, optimized high-performance 3D pulse sequences, and a strategy for resolving ambiguities in distance restraint extraction, has a broad applicability to the structure investigation of membrane proteins within their native membranes. Proton dilution achieved through RAP, together with moderate MAS frequencies (40–60 kHz) and efficient, robust pulse sequences, ensures the method’s versatility for studying a wide range of membrane proteins using commercially available ssNMR hardware. Of course, the RAP deuteration scheme, 3D hNHH and hCCH experiments and modeling-assisted approach for revolving assignment ambiguity established in this study, can also be used to simplified in vitro system, such as membrane proteins reconstituted in lipid bilayers, as well as other solid protein samples^13,41,74,75^. When combined with cryogenic probes^76,77^, dynamic nuclear polarization (DNP)^28,78,79^ and ultrahigh-field NMR^73,80^, this methodology offers enhanced capabilities for the structural analysis of membrane proteins in native membrane, thereby deepening our understanding of fundamental biological processes in native cellular environments and accelerating structure-based drug discovery^13^.

## CONCLUSION

In this study, we determined high resolution structures of MscL in native cellular membranes by newly established ^1^H-detected ssNMR method. We developed a comprehensive *in situ* ssNMR approach comprising three integral components: a partial deuteration protocol for cellular membrane sample preparation, high sensitivity 3D experiments, and a strategy for resolving assignment ambiguities. By enhancing target protein signals while suppressing background protein content, we established an effective sample preparation protocol for partially deuterated proteins. At moderate MAS rate (40-60 kHz), the MscL sample prepared using this protocol produced high quality ^1^H ssNMR spectra suitable for structure determination, with background protein signals completely suppressed in ^1^H-^15^N and ^1^H-^13^C correlation spectra. The RAP technique employed allows partial protonation of all residues, including those in transmembrane domain, and permits different protonation levels for amide and side-chain protons to meet different experimental requirements. For collection of ^1^H^N^-^1^H^N^ and ^1^H^N^-^1^H^SC^ distance restraints, we prepared RAP samples with ∼ 25% for amide protons to maintain adequate sensitivity and ∼10% for side-chain protons to optimize spectral resolution. This optimization reduces side-chain proton linewidths to ∼ 50 Hz, significantly facilitates the extraction of critical ^1^H^N^-^1^H^SC^ distance constraints. To address challenges in assigning side-chain proton chemical shifts, we acquired 3D hCCH spectra. Sensitivity was maximized using a novel ^13^C-^13^C recoupling sequence, PC4, which enhanced polarization transfer efficiency by approximately 50%. Additionally, for key 3D ^1^H-^1^H correlation spectra (e.g., 3D hNHH), we implemented the band-selective ^1^H-^1^H recoupling sequence MODIST, which offers 50% higher recoupling efficiency than RFDR, thereby reliably capturing weak yet essential ^1^H-^1^H cross-peaks. To resolve assignment ambiguity, we integrated 3D hNHH experimental data with CS-Rosetta structural modeling and extracted 92 long range ^1^H-^1^H distance restraints, including 49 long-range ^1^H^N^-^1^H^SC^ restraints essential for structure convergence. High resolution structures of MscL with a backbone RMSD of 1.9 Å were calculated based on ssNMR experimental restraints. The *in situ* ssNMR structures of MscL resemble those obtained in detergent micelles, with only minor variations in secondary structures and tilt angles of transmembrane helices. Compared to ^13^C detection, ^1^H detected *in situ* ssNMR offers significant advantages in spectral sensitivity, demonstrating great potential for broad application in determining membrane protein structures in native membranes.

## METHODS

### Preparation of [^1^H,^13^C,^15^N]-Labeled MscL Cellular Membrane Samples

The large-conductance mechanosensitive channel gene from *Methanosarcina acetivorans* (MaMscL)^52^ with an N-terminal 6×His tag was expressed in *E. coli* BL21(DE3) cells using the pET-15b plasmid. All cellular samples were prepared following a common initial procedure^53^: cells were grown in M9 media with [U-^2^H,^13^C] glucose and ^15^NH_4_Cl as sole carbon and nitrogen sources, induced with 1 mM IPTG, harvested, and lysed by ultrasonication. Total cellular membranes were pelleted by ultracentrifugation at 58,500 × g for 1 hour. Inner membranes were subsequently purified via sucrose density gradient centrifugation, with the centrifugation speed optimized for each sample’s deuteration level as specified below.

**Sample I** was prepared in 100% D_2_O. The inner membranes were isolated by sucrose gradient centrifugation at 76,209 × g for 16 hours. The harvested membrane fraction was then subjected to back-exchange against 100% H_2_O via ultrasonication to replace labile deuterons. After freeze-drying, the membranes were rehydrated with 100% H_2_O at a water-to-membrane mass ratio of 0.4:1 and packed into a 1.9 mm rotor.

**Sample II** was cultured in media with an isotopic composition of 25% H_2_O/75% D_2_O. The sucrose gradient centrifugation step was adjusted to 65,564 × g for 16 hours to account for the altered buoyant density. Following isolation, the inner membranes were back-exchanged into and rehydrated with 100% H_2_O, maintaining a water-to-membrane mass ratio of 0.4:1.

**Sample III** was grown in a medium containing 10% H_2_O/90% D_2_O. The inner membranes were separated by sucrose gradient centrifugation at 76,209 × g for 16 hours. To preserve the specific protonation environment, the back-exchange step after membrane isolation and the final rehydration of the lyophilized material were both performed using a 10% H_2_O/90% D_2_O mixture.

**Sample IV** was prepared in 25% H_2_O/75% D_2_O media. The sucrose gradient centrifugation was conducted at 65,564 × g for 16 hours, consistent with Sample II. A distinct step for this sample was the use of a 25% H_2_O/75% D_2_O mixture for both the back-exchange process and the final rehydration of the lyophilized inner membranes, ensuring a consistent protonation level throughout the preparation.

### Preparation of MscL liposome sample

The [10% ^1^H, ^13^C, ^15^N]-labeled MscL protein was expressed using the pET-15b plasmid harboring an N-terminal 6×His tag in E. coli C43(DE3) cells. MscL colonies were inoculated into unlabeled LB medium (10%H_2_O/90%D_2_O) supplemented with 100 ug/mL ampicillin and grown at 310 K for 8 hours. A 0.4 mL aliquot of the culture was centrifuged, and the pellet was resuspended in 25 mL of M9 medium (10%H_2_O/90%D_2_O) containing 1 g/L NH_4_Cl and 6 g/L [U-^2^H] D-glucose, followed by overnight incubation at 310 K. The entire 25 mL culture was then used to inoculate 225 mL of M9 medium (10%H_2_O/90%D_2_O) containing 1 g/L NH_4_Cl and 6 g/L [U-^2^H,^13^C] D-glucose. Incubation continued at 310 K until the OD600 reached approximately 1.1. Protein expression was induced by adding 1 mM IPTG, followed by additional incubation for 6 hours at 310 K. Cells were harvested by centrifugation at 6,371 × g for 10 minutes and resuspended in lysis buffer (20 mM Tris, 100 mM NaCl, pH 8.0). The subsequent procedures for protein purification and reconstitution were performed as previously described, with the modification that the purified MscL was reconstituted into liposomes composed of a 4:1 (w/w) mixture of POPC and POPG. The resulting proteoliposome pellets were suspended in a 10%H_2_O/90%D_2_O mixture and subjected to ultrasonication and rotational exchange. Proteoliposomes were collected by ultracentrifugation at 419,832 × g for 3 hours. After freeze-drying, the samples were rehydrated with a 10%H_2_O/90%D_2_O mixture at a water-to-proteoliposome mass ratio of approximately 0.4:1 and packed into a 1.9 mm rotor for solid-state NMR analysis.

### Proton-detected ssNMR experiments

All solid-state NMR (ssNMR) experiments were conducted on a Bruker Avance III 800 MHz spectrometer equipped with a 1.9 mm quadra-resonance (HCND) probe. Spectra were acquired under magic-angle spinning (MAS) at 40 kHz, including 3D ^1^H-detected hCCH and hNHH experiments, as well as a series of 2D ^15^N-^1^H heteronuclear single-quantum correlation (HSQC) spectra. The sample temperature was calibrated using internal H_2_O reference and maintained at 283 K for all experiments. Chemical shifts were referenced to adamantane as an external standard, with the methylene carbon signal set to 40.48 ppm^81^. All NMR data were processed with NMRpipe^82^ and analyzed by Sparky^83^.

The key parameters for the ^1^H-detected 2D hNH experiments are as follows. ^1^H→^15^N cross-polarization (CP)^58^ was performed with a contact time of 1000 us, using radio frequency (RF) field strengths of approximately 70 kHz for ^1^H and 30 kHz for ^15^N to satisfy the Hartmann–Hahn condition. The reverse ^15^N→^1^H CP step employed similar field strengths, with a contact time of 0.2 ms to ensure efficient one-bond polarization transfer. A linear ramp from 80% to 100% was applied on the ^1^H channel during CP. WALTZ-16 decoupling^84^ was applied at a power of 10 kHz (0.25ωr), while XY decoupling was set to 50 kHz. A T delay of 44 ms was used for water suppression^85^. The spoil pulse for water suppression had a duration of 1.2 ms and power matching the first ^1^H CP pulse. The water suppression pulse train was applied at 11 kHz, with pulse durations of 20 ms and 34 ms. Typical 90° pulse lengths were 2.2 μs (^1^H), 4.5 μs (^13^C), and 5.5 μs (^15^N). To limit multi-bond ^15^N-^1^H polarization transfer and avoid interference from folded signals in indirect dimensions, the ^15^N-^1^H CP contact time for Sample I was set to 50 μs, and the ^15^N spectral bandwidth was set to 20 kHz. ssNMR experimental details and parameters are described in **Table S1**.

In the 3D hCCH experiments, ^1^H-^13^C polarization transfer was achieved via CP, followed by ^13^C chemical shift evolution. ^13^C-^13^C transfer was accomplished using the PC4 sequence^59^, with the ^13^C carrier frequency set at 40 ppm and a duration of 1.44 ms. During PC4, the ^1^H decoupling power was 70 kHz, and the ^13^C RF field strength was 5.5 kHz. The final ^13^C-^1^H transfer step employed an INEPT sequence^60^ to selectively transfer polarization to directly bonded protons, enabling the assignment of ^1^H chemical shifts.

In 3D hNHH experiments, ^1^H-^1^H polarization transfer was achieved using RFDR^62^ and MODIST^63,64^ recoupling modules. For RFDR, the ^1^H RF field strength was 100 kHz, with a pulse width of 5 μs and recoupling times of 2 ms and 4 ms. For MODIST, the ^1^H RF field strength was set to 20 kHz (0.5ωr), with a pulse width of 6.25us and recoupling times of 6 ms and 12 ms.

### Structural restraints. Dihedral angle restraints

Dihedral angle restraints were predicted from assigned chemical shifts of N, Cα, C’, and Cβ nuclei using TALOS+^86^. Only predictions classified as ‘good’ by the program were adopted as restraints, with the associated prediction errors used to define the restraint bounds. This filtering resulted in a total of 67 φ/ψ dihedral angle restraints. **Hydrogen-bond restraints:** Intra-helical hydrogen-bond restraints were introduced for residue pairs where both the chemical shift index and TALOS+ predictions indicated helical conformation. A hydrogen bond was defined by the distance between the carbonyl oxygen of residue i (Oi) and the amide nitrogen of residue i+4 (Ni+4), with the distance restraint set between 1.5 and 3.5 Å. A total of 43 hydrogen-bond restraints were incorporated for MscL.

### ^1^H-^1^H distance restraints

^1^H-^1^H distance restraints were derived from 3D hNHH spectra acquired on different samples using various recoupling schemes. Specifically, ^1^H^N^–^1^H^N^ distances were obtained from Sample II using both RFDR (**Figure 3a, blue**) and MODIST (**Figure 3a, green**) recoupling. ^1^H^N^–^1^H^Side-chain^ distances were derived from Sample II (**Figure 3b, blue**) and Sample IV (**Figure 3b, red**) using the RFDR scheme. The peaks corresponding to ^1^H-^1^H distance constraints were assigned based on the following process. 1) The cross peaks were hand-picked in Sparky. The criteria used to judge signals in the 3D hNHH spectra were well-resolved and sufficient signal-to-noise ratio (≥6) in 3D hNHH spectra. 2) The possible assignments of the cross-peaks were obtained. A tolerance of ∼0.1 ppm was set for both w2- and w3-dimensional ^1^H chemical shifts considering the linewidth of the peaks (0.1-0.2 ppm) in the 3D hNHH spectra. A tolerance of ∼0.5 ppm was set for w1 dimensional ^15^N chemical shifts considering the linewidth of the peaks (0.6-1.0 ppm) in the 3D hNHH spectra. This procedure produced a number of ambiguous assignments. 3) The ambiguity in the assignments was eliminated by using the following procedures. i) The ambiguity in the assignment of intramonomeric correlations can be significantly reduced by using the MscL pentamer structure determined by CS-Rosetta calculations using the assigned chemical shifts of MscL in cellular membranes as input. ii) The assignments of intra-monomer or inter-monomer correlations were confirmed by the MscL pentamer structure. By using this procedure, we obtained a total of 43 long-range ^1^H^N^-^1^H^N^ distance restraints, 49 long-range ^1^H^N^-^1^H^SC^ distance restraints, and 186 inter-residue restraints(**Table S5**).

### MscL pentameric structures predicted by CS-Rosetta

The initial MscL pentamer structure was predicted using the standard POMONA/CS-RosettaCM protocol within the CS-Rosetta package^87,88^. A total of 74 backbone (^15^N, ^13^Cα, ^13^C’) and 59 ^13^Cβ chemical shifts were used as input for fragment and template selection^54^. Only fragments and templates from PDB proteins with less than 30% sequence similarity to MscL were used in the subsequent structure generation. Notably, all calculations were performed in a membrane-environment context. From 10,000 all-atom models generated, the 50 lowest-energy structures were selected as the final ensemble of initial pentamer structures for subsequent analysis.

### Assignment of Intra- and Inter-monomer ^1^H–^1^H Distance Restraints

Distance restraints were derived from five 3D hNHH spectra acquired on Samples II and IV using RFDR and MODIST recoupling schemes. Specifically, ^1^H^N^–^1^H^N^ distances were obtained from Sample II with both RFDR and MODIST recoupling, while ^1^H^N^–^1^H^Side-chain^ distances were derived from Sample II and Sample IV using the RFDR scheme.

The assignment of cross-peaks was conducted through a multi-step procedure in Sparky. First, peaks were manually selected based on the criteria of being well-resolved and having a signal-to-noise ratio (SNR) ≥ 6, with typical linewidths of 0.1–0.2 ppm in the ^1^H dimensions. For the assignment of potential correlations, chemical shift tolerances were set to ∼0.1 ppm for both w2 and w3 (^1^H) dimensions, and ∼0.5 ppm for the w1 (^15^N) dimension, consistent with the observed spectral linewidths.

To resolve the inherent ambiguity in the initial assignments, the CS-Rosetta-predicted pentamer structures of MscL were employed. At this stage, both intra- and inter-monomer assignments were considered as possibilities. A correlation was only accepted if the distance between the candidate atoms was within 9.0 Å in the pentameric template. The setting of this upper limit takes into account both the distance constraint derived from the 4.0 ms RFDR mixing time (6-8 Å)^48,69^ and the uncertainty of the initial structural model generated by CS-Rosetta.

This rigorous process ultimately yielded two definitive sets of constraints: 49 unambiguous long-range intramonomer distance restraints, and 32 unambiguous and 11 ambiguous long-range intermonomer distance restraints. All long-range distance restraints are summarized in **Table S4**.

### MscL pentameric structures calculated by Xplor-NIH

The final MscL pentamer structures were calculated using Xplor-NIH (version 2.47)^89^. The calculation incorporated standard terms for bond lengths, bond angles, and improper angles to enforce correct covalent geometry, along with statistical torsion angle potentials and a gyration volume term. The hydrogen bond database potential (HBPot) was used to improve hydrogen bond geometry.

Folding calculations for the full-length, single-chain MscL protomer were initiated from extended chains. A total of 1000 structures were calculated using molecular dynamics simulated annealing in torsion angle space, followed by final gradient minimization in Cartesian space. The experimental restraints included 319 unique non-redundant ^1^H–^1^H distances, 43 hydrogen bonds, and 67 TALOS+-derived torsion angle restraints per MscL chain.

The simulated annealing protocol began with constant-temperature molecular dynamics at 3500 K, run for the shorter of 800 ps or 10,000 steps. The temperature was then reduced in steps of 25 K. At each temperature, dynamics were run for the shorter of 2 ps or 500 steps. The force constant for distance restraints was ramped from 5 to 50 kcal mol^−1^ Å^−2^. Dihedral angle restraints were disabled at 3500 K but enabled during annealing with a force constant of 200 kcal mol^−1^ rad^−2^. The force constant for the gyration term was geometrically increased from 0.002 to 1. After simulated annealing, the structures were subjected to final energy minimization using Powell’s method.

This procedure yielded an ensemble of 1000 structures. The 10 lowest-energy structures exhibited a backbone atom RMSD of 1.9 ± 0.4 Å. Structure quality was assessed using PROCHECK^90^ and MolProbity^91^, with the corresponding statistics provided in **Table S7**.

## Supporting information

SI_MaMscL_Structures

## ASSOCIATED CONTENT

This material is available free of charge via the Internet at http://pubs.acs.org.

## Author Contributions

\# H.X. and W.J.Z. contributed equally to this work

## Notes

J.Y. designed research; H.X., H.T. and W.J.Z. conducted experiments; H.X. and H.T. analyzed data; all authors wrote the manuscript.

## ACKNOWLEDGMENT

This work was funded by grants from the National Natural Science Foundation of China (22434003, 22577099, 22507127, 21927801, 21921004 and 22004124), the Chinese Academy of Sciences (XDB0540000), the Hubei Provincial Natural Science Foundation of China (2024AFA005 and 2025AFB512), and the China Postdoctoral Science Foundation (2024M752494, GZC20241290 and 2020M672455).

## Notes

### Competing Interest Statement

The authors have declared no competing interest.

### Summary of Updates

Completed the author list and funding agencies.

## REFERENCES

(1) Zhou, H.-X.; Cross, T. A. Influences of Membrane Mimetic Environments on Membrane Protein Structures. Annu. Rev. Biophys. 2013, 42 (1), 361–392.

(2) Cross, T. A.; Sharma, M.; Yi, M.; Zhou, H.-X. Influence of Solubilizing Environments on Membrane Protein Structures. Trends Biochem. Sci. 2011, 36 (2), 117–125.

(3) Enkavi, G.; Javanainen, M.; Kulig, W.; Róg, T.; Vattulainen, Multiscale Simulations of Biological Membranes: The Challenge to Understand Biological Phenomena in a Living Substance. Chem. Rev. 2019, 119 (9), 5607–5774.

(4) Choy, B. C.; Cater, R. J.; Mancia, F.; Pryor, E. E. A 10-Year Meta-Analysis of Membrane Protein Structural Biology: Detergents, Membrane Mimetics, and Structure Determination Techniques. Biochim. Biophys. Acta (BBA) - Biomembr. 2021, 1863 (3), 183533.

(5) Autzen, H. E.; Julius, D.; Cheng, Y. Membrane Mimetic Systems in CryoEM: Keeping Membrane Proteins in Their Native Environment. Curr. Opin. Struct. Biol. 2019, 58, 259–268.

(6) Nygaard, R.; Kim, J.; Mancia, F. Cryo-Electron Microscopy Analysis of Small Membrane Proteins. Curr. Opin. Struct. Biol. 2020, 64, 26–33.

(7) Danmaliki, G. I.; Hwang, P. M. Solution NMR Spectroscopy of Membrane Proteins. Biochim. Biophys. Acta (BBA) - Biomembr. 2020, 1862 (9), 183356.

(8) Baldus, M. A Solid View of Membrane Proteins In Situ. Biophys. J. 2015, 108 (7), 1585–1586.

(9) Cross, T. A. YidC: Evaluating the Importance of the Native Environment. Structure 2018, 26 (1), 2–4.

(10) Xie, H.; Gan, Y.; Zhao, W.; Duan, M.; Shen, Y.; Tan, H.; Tong, Q.; Yongxiang Zhao; Yang, J. Dynamic Structures of a Membrane Transporter in Native Cellular Membranes. Sci. Adv. 2025, eadv4583.

(11) Xie, H.; Zhao, Y.; Zhao, W.; Chen, Y.; Liu, M.; Yang, J. Solid-State NMR Structure Determination of a Membrane Protein in E. Coli Cellular Inner Membrane. Sci. Adv. 2023, 9 (44), eadh4168.

(12) Zhao, Y.; Xie, H.; Wang, L.; Shen, Y.; Chen, W.; Song, B.; Zhang, Z.; Zheng, A.; Lin, Q.; Fu, R.; Wang, J.; Yang, J. Gating Mechanism of Aquaporin Z in Synthetic Bilayers and Native Membranes Revealed by Solid-State NMR Spectroscopy. J. Am. Chem. Soc. 2018, 140 (25), 7885–7895.

(13) Marassi, F. M.; Pintacuda, G. Solid-State NMR of Membrane Proteins in Situ. Curr. Opin. Struct. Biol. 2025, 94, 103129.

(14) Narasimhan, S.; Pinto, C.; Lucini Paioni, A.; Van Der Zwan, J.; Folkers, G. E.; Baldus, M. Characterizing Proteins in a Native Bacterial Environment Using Solid-State NMR Spectroscopy. Nat. Protoc. 2021, 16 (2), 893–918.

(15) Narasimhan, S.; Folkers, G. E.; Baldus, M. When Small Becomes Too Big: Expanding the Use of In-cell Solid-state NMR Spectroscopy. ChemPlusChem 2020, 85 (4), 760–768.

(16) Brown, L. S.; Ladizhansky, V. Membrane Proteins in Their Native Habitat as Seen by Solid-state NMR Spectroscopy. Protein Sci. 2015, 24 (9), 1333–1346.

(17) Renault, M.; Tommassen-van Boxtel, R.; Bos, M. P.; Post, J. A.; Tommassen, J.; Baldus, M. Cellular Solid-State Nuclear Magnetic Resonance Spectroscopy. Proc. Natl. Acad. Sci. 2012, 109 (13), 4863–4868.

(18) Marley, J.; Lu, M.; Bracken, C. A Method for Efficient Isotopic Labeling of Recombinant Proteins. J. Biomol. NMR 2001, 20 (1), 71–75.

(19) Baker, L. A.; Daniëls, M.; Van Der Cruijsen, E. A. W.; Folkers, G. E.; Baldus, M. Efficient Cellular Solid-State NMR of Membrane Proteins by Targeted Protein Labeling. J. Biomol. NMR 2015, 62 (2), 199–208.

(20) Baker, L. A.; Sinnige, T.; Schellenberger, P.; De Keyzer, J.; Siebert, C. A.; Driessen, A. J. M.; Baldus, M.; Grünewald, K. Combined 1H-Detected Solid-State NMR Spectroscopy and Electron Cryotomography to Study Membrane Proteins across Resolutions in Native Environments. Structure 2018, 26 (1), 161-170.e3.

(21) Ward, M. E.; Wang, S.; Munro, R.; Ritz, E.; Hung, I.; Gor’kov, P. L.; Jiang, Y.; Liang, H.; Brown, L. S.; Ladizhansky, V. In Situ Structural Studies of Anabaena Sensory Rhodopsin in the E. Coli Membrane. Biophys. J. 2015, 108 (7), 1683–1696.

(22) Fu, R.; Wang, X.; Li, C.; Santiago-Miranda, A. N.; Pielak, G. J.; Tian, F. In Situ Structural Characterization of a Recombinant Protein in Native Escherichia Coli Membranes with Solid-State Magic-Angle-Spinning NMR. J. Am. Chem. Soc. 2011, 133 (32), 12370–12373.

(23) Higman, V. A.; Varga, K.; Aslimovska, L.; Judge, P. J.; Sperling, L. J.; Rienstra, C. M.; Watts, A. The Conformation of Bacteriorhodopsin Loops in Purple Membranes Resolved by Solid-state MAS NMR Spectroscopy. Angew. Chem. Int. Ed. 2011, 50 (36), 8432–8435.

(24) Jacso, T.; Franks, W. T.; Rose, H.; Fink, U.; Broecker, J.; Keller, S.; Oschkinat, H.; Reif, B. Characterization of Membrane Proteins in Isolated Native Cellular Membranes by Dynamic Nuclear Polarization Solid-state NMR Spectroscopy without Purification and Reconstitution. Angew. Chem. Int. Ed. 2012, 51 (2), 432–435.

(25) Kulminskaya, N. V.; Pedersen, M.Ø.; Bjerring, M.; Underhaug, J.; Miller, M.; Frigaard, N.; Nielsen, J. T.; Nielsen, N. Chr. In Situ Solid-state NMR Spectroscopy of Protein in Heterogeneous Membranes: The Baseplate Antenna Complex of Chlorobaculum Tepidum. Angew. Chem. Int. Ed. 2012, 51 (28), 6891–6895.

(26) Miao, Y.; Qin, H.; Fu, R.; Sharma, M.; Can, T. V.; Hung, I.; Luca, S.; Gor’kov, P. L.; Brey, W. W.; Cross, T. A. M2 Proton Channel Structural Validation from Full-length Protein Samples in Synthetic Bilayers and E. Coli Membranes. Angew. Chem. Int. Ed. 2012, 51 (33), 8383–8386.

(27) Renault, M.; Pawsey, S.; Bos, M. P.; Koers, E. J.; Nand, D.; Tommassen-van Boxtel, R.; Rosay, M.; Tommassen, J.; Maas, W. E.; Baldus, M. Solid-state NMR Spectroscopy on Cellular Preparations Enhanced by Dynamic Nuclear Polarization. Angew. Chem. Int. Ed. 2012, 51 (12), 2998–3001.

(28) Kaplan, M. Probing a Cell-Embedded Megadalton Protein Complex by DNP-Supported Solid-State NMR. Nat. Methods 2015.

(29) Shahid, S. A.; Nagaraj, M.; Chauhan, N.; Franks, T. W.; Bardiaux, B.; Habeck, M.; Orwick-Rydmark, M.; Linke, D.; van Rossum, B. Solid-state NMR Study of the YadA Membrane-anchor Domain in the Bacterial Outer Membrane. Angew. Chem. Int. Ed. 2015, 54 (43), 12602–12606.

(30) Kaplan, M.; Narasimhan, S.; De Heus, C.; Mance, D.; Van Doorn, S.; Houben, K.; Popov-čeleketić, D.; Damman, R.; Katrukha, E. A.; Jain, P.; Geerts, W. J. C.; Heck, A. J. R.; Folkers, G. E.; Kapitein, L. C.; Lemeer, S.; Van Bergen En Henegouwen, P.M.P.; Baldus, M. EGFR Dynamics Change during Activation in Native Membranes as Revealed by NMR. Cell 2016, 167 (5), 1241-1251.e11.

(31) Medeiros-Silva, J.; Mance, D.; Daniëls, M.; Jekhmane, S.; Houben, K.; Baldus, M.; Weingarth, M. 1 H-detected Solid-state NMR Studies of Water-inaccessible Proteins In Vitro and In Situ. Angew. Chem. Int. Ed. 2016, 55 (43), 13606–13610.

(32) Pinto, C.; Mance, D.; Julien, M.; Daniels, M.; Weingarth, M.; Baldus, M. Studying Assembly of the BAM Complex in Native Membranes by Cellular Solid-State NMR Spectroscopy. J. Struct. Biol. 2019, 206 (1), 1–11.

(33) Azadi-Chegeni, F.; Ward, M. E.; Perin, G.; Simionato, D.; Morosinotto, T.; Baldus, M.; Pandit, A. Conformational Dynamics of Light-Harvesting Complex II in a Native Membrane Environment. Biophysical Journal 2021, 120 (2), 270–283.

(34) Zhang, Y.; Gan, Y.; Zhao, W.; Zhang, X.; Zhao, Y.; Xie, H.; Yang, J. Membrane Protein Structures in Native Cellular Membranes Revealed by Solid-State NMR Spectroscopy. JACS Au 2023, 3 (12), 3412–3423.

(35) Le Marchand, T.; Schubeis, T.; Bonaccorsi, M.; Paluch, P.; Lalli, D.; Pell, A. J.; Andreas, L. B.; Jaudzems, K.; Stanek, J.; Pintacuda, G. ^1^ H-Detected Biomolecular NMR under Fast Magic-Angle Spinning. Chem. Rev. 2022, 122 (10), 9943–10018.

(36) Mandala, V. S.; Hong, M. High-Sensitivity Protein Solid-State NMR Spectroscopy. Curr. Opin. Struct. Biol. 2019, 58, 183–190.

(37) Andreas, L. B.; Jaudzems, K.; Stanek, J.; Lalli, D.; Bertarello, A.; Le Marchand, T.; Cala-De Paepe, D.; Kotelovica, S.; Akopjana, I.; Knott, B.; Wegner, S.; Engelke, F.; Lesage, A.; Emsley, L.; Tars, K.; Herrmann, T.; Pintacuda, G. Structure of Fully Protonated Proteins by Proton-Detected Magic-Angle Spinning NMR. Proc. Natl. Acad. Sci. 2016, 113 (33), 9187–9192.

(38) Schubeis, T.; Le Marchand, T.; Andreas, L. B.; Pintacuda, G. 1H Magic-Angle Spinning NMR Evolves as a Powerful New Tool for Membrane Proteins. J. Magn. Reson. 2018, 287, 140–152.

(39) Shcherbakov, A. A.; Medeiros-Silva, J.; Tran, N.; Gelenter, M. D.; Hong, M. From Angstroms to Nanometers: Measuring Interatomic Distances by Solid-State NMR. Chem. Rev. 2022, 122 (10), 9848–9879.

(40) Reif, B. Deuteration for High-Resolution Detection of Protons in Protein Magic Angle Spinning (MAS) Solid-State NMR. Chem. Rev. 2022, 122 (10), 10019–10035.

(41) Hu, S.; Li, J.; Yuan, F.; Zhang, J.; Cheng, X.; Xiang, S.; Tian, C.; Gong, W.; Liu, T.; Shi, C. Structural Mechanism of Insect Cuticular Protein Binding to Chitin Revealed by Solid-State NMR. J. Am. Chem. Soc. 2025.

(42) Lalli, D.; Idso, M. N.; Andreas, L. B.; Hussain, S.; Baxter, N.; Han, S.; Chmelka, B. F.; Pintacuda, G. Proton-Based Structural Analysis of a Heptahelical Transmembrane Protein in Lipid Bilayers. J. Am. Chem. Soc. 2017, 139 (37), 13006–13012.

(43) Linser, R.; Bardiaux, B.; Andreas, L. B.; Hyberts, S. G.; Morris, V. K.; Pintacuda, G.; Sunde, M.; Kwan, A. H.; Wagner, G. Solid-State NMR Structure Determination from Diagonal-Compensated, Sparsely Nonuniform-Sampled 4D Proton–Proton Restraints. J. Am. Chem. Soc. 2014, 136 (31), 11002–11010.

(44) Zhou, D. H.; Shea, J. J.; Nieuwkoop, A. J.; Franks, W. T.; Wylie, B. J.; Mullen, C.; Sandoz, D.; Rienstra, C. M. Solid-state Protein-structure Determination with Proton-detected Triple-resonance 3D Magic-angle-spinning NMR Spectroscopy. Angew. Chem. Int. Ed. 2007, 46 (44), 8380–8383.

(45) Shcherbakov, A. A.; Hisao, G.; Mandala, V. S.; Thomas, N. E.; Soltani, M.; Salter, E. A.; Davis, J. H.; Henzler-Wildman, K. A.; Hong, M. Structure and Dynamics of the Drug-Bound Bacterial Transporter EmrE in Lipid Bilayers. Nat. Commun. 2021, 12 (1), 172.

(46) Najbauer, E. E.; Tekwani Movellan, K.; Giller, K.; Benz, R.; Becker, S.; Griesinger, C.; Andreas, L. B. Structure and Gating Behavior of the Human Integral Membrane Protein VDAC1 in a Lipid Bilayer. J. Am. Chem. Soc. 2022, 144 (7), 2953–2967.

(47) Andreas, L. B.; Reese, M.; Eddy, M. T.; Gelev, V.; Ni, Q. Z.; Miller, E. A.; Emsley, L.; Pintacuda, G.; Chou, J. J.; Griffin, R. G. Structure and Mechanism of the Influenza a M2_18–60_Dimer of Dimers. J. Am. Chem. Soc. 2015, 137 (47), 14877–14886.

(48) Linser, R.; Bardiaux, B.; Higman, V.; Fink, U.; Reif, B. Structure Calculation from Unambiguous Long-Range Amide and Methyl^1^ H−^1^ H Distance Restraints for a Microcrystalline Protein with MAS Solid-State NMR Spectroscopy. J. Am. Chem. Soc. 2011, 133 (15), 5905–5912.

(49) Retel, J. S.; Nieuwkoop, A. J.; Hiller, M.; Higman, V. A.; Barbet-Massin, E.; Stanek, J.; Andreas, L. B.; Franks, W. T.; Van Rossum, B.-J.; Vinothkumar, K. R.; Handel, L.; De Palma, G. G.; Bardiaux, B.; Pintacuda, G.; Emsley, L.; Kühlbrandt, W.; Oschkinat, H. Structure of Outer Membrane Protein G in Lipid Bilayers. Nat. Commun. 2017, 8 (1), 2073.

(50) Cox, C. D.; Bavi, N.; Martinac, B. Bacterial Mechanosensors. Annu. Rev. Physiol. 2018, 80 (1), 71–93.

(51) Sukharev, S. I.; Blount, P.; Martinac, B.; Kung; Ching. MECHANOSENSITIVE CHANNELS OFESCHERICHIA COLI:The MscL Gene, Protein, and Activities. Annual Review of Physiology, 1997, 59, 633–657.

(52) Li, J.; Guo, J.; Ou, X.; Zhang, M.; Li, Y.; Liu, Z. Mechanical Coupling of the Multiple Structural Elements of the Large-Conductance Mechanosensitive Channel during Expansion. Proc. Natl. Acad. Sci. 2015, 112 (34), 10726–10731.

(53) Tan, H.; Zhao, W.; Duan, M.; Zhao, Y.; Zhang, Y.; Xie, H.; Tong, Q.; Yang, J. Native Cellular Membranes Facilitate Channel Activity of MscL by Enhancing Slow Collective Motions of Its Transmembrane Helices. J. Am. Chem. Soc. 2024, 146 (46), 31472–31485.

(54) Zhang, X.; Zhang, Y.; Tang, S.; Ma, S.; Shen, Y.; Chen, Y.; Tong, Q.; Li, Y.; Yang, J. Hydrophobic Gate of Mechanosensitive Channel of Large Conductance in Lipid Bilayers Revealed by Solid-State NMR Spectroscopy. J. Phys. Chem. B 2021, 125 (10), 2477–2490.

(55) Asami, S.; Schmieder, P.; Reif, B. High Resolution^1^ H-Detected Solid-State NMR Spectroscopy of Protein Aliphatic Resonances: Access to Tertiary Structure Information. J. Am. Chem. Soc. 2010, 132 (43), 15133–15135.

(56) Asami, S.; Szekely, K.; Schanda, P.; Meier, B. H.; Reif, B. Optimal Degree of Protonation for 1H Detection of Aliphatic Sites in Randomly Deuterated Proteins as a Function of the MAS Frequency. J Biomol NMR 2012, 54 (2), 155–168.

(57) Asami, S.; Reif, B. Proton-Detected Solid-State NMR Spectroscopy at Aliphatic Sites: Application to Crystalline Systems. Acc. Chem. Res. 2013, 46 (9), 2089–2097.

(58) Baldus, M.; Petkova, A. T.; Herzfeld, J.; Griffin, R. G. Cross Polarization in the Tilted Frame: Assignment and Spectral Simplification in Heteronuclear Spin Systems. Mol, Phys, 1998, 95 (6), 1197–1207.

(59) Xiao, H.; Zhao, W.; Zhang, Y.; Kang, H.; Zhang, Z.; Yang, J. Selective Correlations between Aliphatic 13C Nuclei in Protein Solid-State NMR. J. Magn. Reson. 2024, 365, 107730.

(60) Morris, G. A.; Freeman, R. Enhancement of Nuclear Magnetic Resonance Signals by Polarization Transfer. J. Am. Chem. Soc. 1979, 101 (3), 760–762.

(61) Russell, R. W.; Fritz, M. P.; Kraus, J.; Quinn, C. M.; Polenova, T.; Gronenborn, A. M. Accuracy and Precision of Protein Structures Determined by Magic Angle Spinning NMR Spectroscopy: For Some “with a Little Help from a Friend.” J. Biomol. NMR 2019, 73 (6–7), 333–346.

(62) Bennett, A. E.; Rienstra, C. M.; Griffiths, J. M.; Zhen, W.; Lansbury, Jr., Peter T.; Griffin, R. G. Homonuclear Radio Frequency-Driven Recoupling in Rotating Solids. J. Chem. Phys. 1998, 108 (22), 9463–9479.

(63) Nimerovsky, E.; Stampolaki, M.; Varkey, A. C.; Becker, S.; Andreas, L. B. Analysis of the MODIST Sequence for Selective Proton–Proton Recoupling. J. Phys. Chem. A 2025, 129 (1), 317–329.

(64) Nimerovsky, E.; Najbauer, E. E.; Movellan, K. T.; Xue, K.; Becker, S.; Andreas, L. B. Modest Offset Difference Internuclear Selective Transfer via Homonuclear Dipolar Coupling. J. Phys. Chem. Lett. 2022, 13 (6), 1540–1546.

(65) Haswell, E. S.; Phillips, R.; Rees, D. C. Mechanosensitive Channels: What Can They Do and How Do They Do It? Structure 2011, 19 (10), 1356–1369.

(66) Mandala, V. S.; McKay, M. J.; Shcherbakov, A. A.; Dregni, J.; Kolocouris, A.; Hong, M. Structure and Drug Binding of the SARS-CoV-2 Envelope Protein Transmembrane Domain in Lipid Bilayers. Nat. Struct. Mol. Biol. 2020, 27 (12), 1202–1208.

(67) Loquet, A.; Habenstein, B.; Lange, A. Structural Investigations of Molecular Machines by Solid-State NMR. Acc. Chem. Res. 2013, 46 (9), 2070–2079.

(68) Mandala, V. S.; Loftis, A. R.; Shcherbakov, A. A.; Pentelute, L.; Hong, M. Atomic Structures of Closed and Open Influenza B M2 Proton Channel Reveal the Conduction Mechanism. Nat. Struct. Mol. Biol. 2020, 27 (2), 160–167.

(69) Agarwal, V.; Penzel, S.; Szekely, K.; Cadalbert, R.; Testori, E.; Oss, A.; Past, J.; Samoson, A.; Ernst, M.; Böckmann, A.; Meier, B. H. De Novo 3D Structure Determination from Sub-milligram Protein Samples by Solid-state 100 kHz MAS NMR Spectroscopy. Angew Chem Int Ed 2014, 53 (45), 12253–12256.

(70) Xie, H.; Zhao, Y.; Wang, J.; Zhang, Z.; Yang, J. Solid-State NMR Chemical Shift Assignments of Aquaporin Z in Lipid Bilayers. Biomol. NMR Assignments 2018, 12 (2), 323–328.

(71) Nishiyama, Y.; Hou, G.; Agarwal, V.; Su, Y.; Ramamoorthy, Ultrafast Magic Angle Spinning Solid-State NMR Spectroscopy: Advances in Methodology and Applications. Chem. Rev. 2023, 123 (3), 918–988.

(72) Sun, Z.; Ollier, C.; Rancz, A.; Thienpont, B.; Grohe, K.; Becker, L.; Purea, A.; Engelke, F.; Wegner, S.; Sturgis, J.; Polenova, T.; Le Marchand, T.; Pintacuda, G. Pushing the Boundaries of Resolution in Solid-State Nuclear Magnetic Resonance of Biomolecules with 160 kHz Magic-Angle Spinning. J. Am. Chem. Soc. 2025, 147 (23), 19433–19437.

(73) Callon, M.; Luder, D.; Malär, A. A.; Wiegand, T.; Římal, V.; Lecoq, L.; Böckmann, A.; Samoson, A.; Meier, B. H. High and Fast: NMR Protein–Proton Side-Chain Assignments at 160 kHz and 1.2 GHz. Chem. Sci. 2023, 14 (39), 10824–10834.

(74) Yarava, J. R.; Gautam, I.; Jacob, A.; Fu, R.; Wang, T. Proton-Detected Solid-State NMR for Deciphering Structural Polymorphism and Dynamic Heterogeneity of Cellular Carbohydrates in Pathogenic Fungi. Biochemistry March 13, 2025.

(75) Daskalov, A.; Martinez, D.; Coustou, V.; El Mammeri, N.; Berbon, M.; Andreas, L. B.; Bardiaux, B.; Stanek, J.; Noubhani, A.; Kauffmann, B.; Wall, J. S.; Pintacuda, G.; Saupe, S. J.; Habenstein, B.; Loquet, A. Structural and Molecular Basis of Cross-Seeding Barriers in Amyloids. Proc. Natl. Acad. Sci. 2021, 118 (1).

(76) Hassan, A.; Quinn, C. M.; Struppe, J.; Sergeyev, I. V.; Zhang, C.; Guo, C.; Runge, B.; Theint, T.; Dao, H. H.; Jaroniec, C. P.; Berbon, M.; Lends, A.; Habenstein, B.; Loquet, A.; Kuemmerle, R.; Perrone, B.; Gronenborn, A. M.; Polenova, T. Sensitivity Boosts by the CPMAS CryoProbe for Challenging Biological Assemblies. J. Magn. Reson. 2020, 311, 106680.

(77) Sučec, I.; Xia, B.; Somberg, N. H.; Wang, Y.; Jo, H.; Li, S.; Perrone, B.; Gao, Z.; Hong, M. Ion Channel Structure and Function of the MERS Coronavirus E Protein. Sci. Adv. 2025, 11 (28), eadx1788.

(78) Biedenbänder, T.; Aladin, V.; Saeidpour, S.; Corzilius, B. Dynamic Nuclear Polarization for Sensitivity Enhancement in Biomolecular Solid-State NMR. Chem. Rev. 2022, 122 (10), 9738–9794.

(79) Frederick, K. K.; Michaelis, V. K.; Corzilius, B.; Ong, T. C.; Jacavone, A. C.; Griffin, R. G.; Lindquist, S. Sensitivity-Enhanced NMR Reveals Alterations in Protein Structure by Cellular Milieus. Cell 2015, 163 (3), 620–628.

(80) Wang, S.; Ravula, T.; Stringer, J. A.; Gor’kov, P. L.; Warmuth, O. A.; Williams, C. G.; Thome, A. F.; Mueller, L. J.; Rienstra, C. M. Ultrahigh-Resolution Solid-State NMR for High–Molecular Weight Proteins on GHz-Class Spectrometers. Sci. Adv. 11 (30), eadx6016.

(81) Morcombe, C. R.; Zilm, K. W. Chemical Shift Referencing in MAS Solid State NMR. J. Magn. Reson. 2003, 162 (2), 479–486.

(82) Delaglio, F.; Grzesiek, S.; Vuister, G. W.; Zhu, G.; Pfeifer, J.; Bax, A. NMRPipe: A Multidimensional Spectral Processing System Based on UNIX Pipes. J. Biomol, NMR. 1995, 6 (3), 277–293.

(83) D. Goddard and D. G. Kneller, “SPARKY 3,” University of California, San Francisco, 2000.

(84) Shaka, A. J.; Keeler, J.; Frenkiel, T.; Freeman, R. An Improved Sequence for Broadband Decoupling: WALTZ-16. J. Magn. Reson. (1969) 1983, 52 (2), 335–338.

(85) Fricke, P.; Chevelkov, V.; Zinke, M.; Giller, K.; Becker, S.; Lange, A. Backbone Assignment of Perdeuterated Proteins by Solid-State NMR Using Proton Detection and Ultrafast Magic-Angle Spinning. Nat. Protoc. 2017, 12 (4), 764–782.

(86) Shen, Y.; Delaglio, F.; Cornilescu, G.; Bax, A. TALOS+: A Hybrid Method for Predicting Protein Backbone Torsion Angles from NMR Chemical Shifts. J Biomol NMR 2009, 44 (4), 213–223.

(87) Shen, Y.; Lange, O.; Delaglio, F.; Rossi, P.; Aramini, J. M.; Liu, G.; Eletsky, A.; Wu, Y.; Singarapu, K. K.; Lemak, A.; Ignatchenko, A.; Arrowsmith, C. H.; Szyperski, T.; Montelione, G. T.; Baker, D.; Bax, A. Consistent Blind Protein Structure Generation from NMR Chemical Shift Data. Proc. Natl. Acad. Sci. 2008, 105 (12), 4685–4690.

(88) Shen, Y.; Bax, A. Homology Modeling of Larger Proteins Guided by Chemical Shifts. Nat, Methods 2015, 12 (8), 747–750.

(89) Schwieters, C. D.; Kuszewski, J. J.; Tjandra, N.; Marius Clore, G. The Xplor-NIH NMR Molecular Structure Determination Package. Journal of Magnetic Resonance 2003, 160 (1), 65–73.

(90) Laskowski, R. A.; MacArthur, M. W.; Moss, D. S.; Thornton, J. M. PROCHECK: A Program to Check the Stereochemical Quality of Protein Structures. J. Appl, Crystallogr, 1993, 26 (2), 283–291.

(91) Chen, V. B.; Arendall, W. B.; Headd, J. J.; Keedy, D. A.; Immormino, R. M.; Kapral, G. J.; Murray, L. W.; Richardson, J. S.; Richardson, D. C. MolProbity: All-Atom Structure Validation for Macromolecular Crystallography. Acta, Crystallogr, D. Biol, Crystallogr, 2010, 66 (Pt 1), 12–21.

