## Supplementary material for "*In Situ* Structure Determination of a Membrane Protein in Native Cellular Membranes by Proton-Detected Solid-State NMR": SI_MaMscL_Structures

**This PDF file includes:**

Figures. S1 to S11

Tables. S1 to S6

References. 1-17

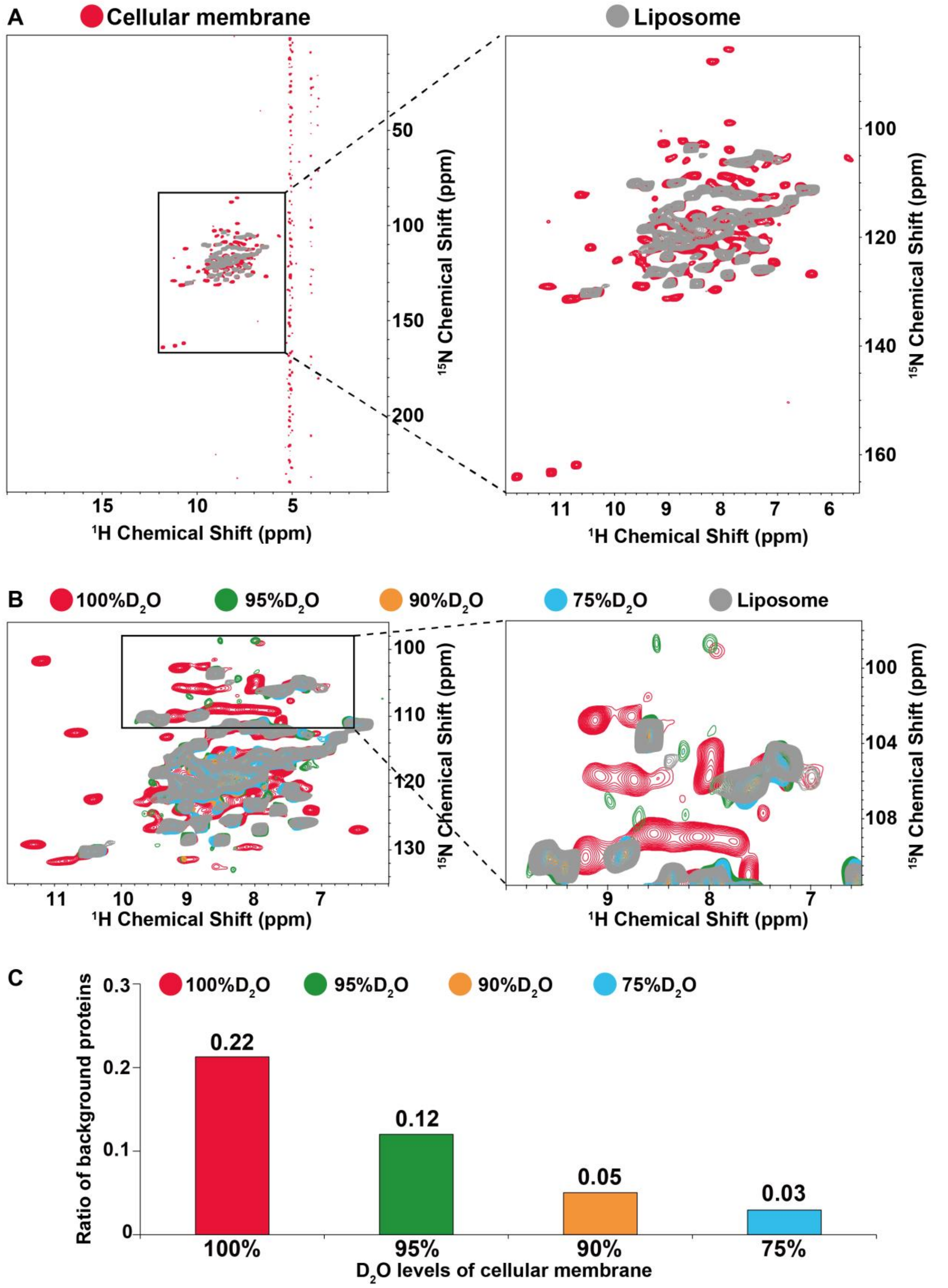


**Figure S1. Characterization of background protein signals in cellular membrane samples prepared in deuterated expression media containing different levels of $D_2O$.** (A) Confirmation of background protein signals in the MscL cellular membrane sample prepared by expressing proteins in media containing 100% $D_2O$. The superimposition of 2D $^{15}N$-$^{1}H$ CP-HSQC spectra of MscL in native membrane samples prepared using the 100% deuterated RAP protocol with H/D back-exchange in 100% $H_2O$ (red) and that of [10% $^{1}H$, $^{13}C$, $^{15}N$] labeled MscL in proteoliposomes (gray). The right panel shows a zoomed-in region of the left panel. To limit the spatial transfer of polarization through multiple $^{15}N$-$^{1}H$ bonds, the $^{15}N$-$^{1}H$ cross-polarization contact time was set to 50 μs in both HSQC spectra. To avoid interference from aliased signals in the indirect dimension on the target signals, a sufficiently wide spectral width (250 ppm) was set for the indirect $^{15}N$ dimension. (B) Background protein signals present in 2D $^{15}N$-$^{1}H$ CP-HSQC of MscL in cellular membrane samples with different deuteration ratios in RAP. The superimposition of 2D $^{15}N$-$^{1}H$ HSQC spectra of MscL in cellular membrane prepared using four RAP cultivation protocols with different deuteration ratios (100% $D_2O$: red; 95% $D_2O$: green; 90% $D_2O$: orange; 75% $D_2O$: cyan) and that of [10% $^{1}H$, $^{13}C$, $^{15}N$] labeled MscL in proteoliposomes (gray). The right panel shows a zoomed-in region of the left panel. (C) The relative contents of background protein signals in 2D $^{15}N$-$^{1}H$ HSQC spectra of MscL as the function of deuteration ratio in RAP in sample preparations.

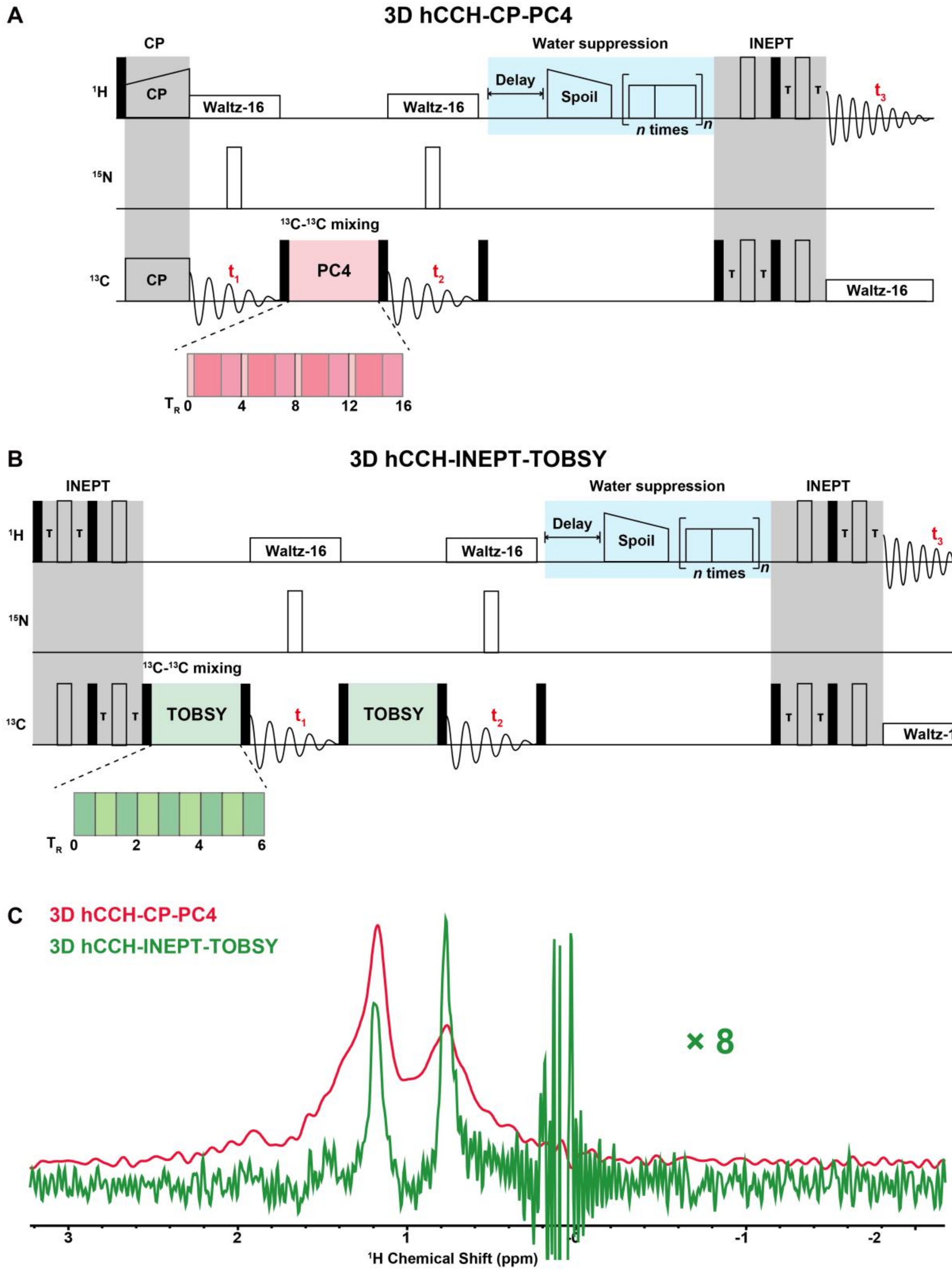


**Figure S2. The 3D hCCH experiment for assignments of methyl proton chemical shift of MscL in native membrane.** (A) Pulse sequence of 3D hCCH-CP-PC4 was used in 3D hCCH experimtns, in which open and filled bars represent 90° and 180° pulses, respectively. In this experiment, $^1$H-$^{13}$C polarization transfer was achieved via cross-polarization (CP)[1], followed by $^{13}$C chemical shift evolution. $^{13}$C-$^{13}$C transfer was accomplished using the PC4 sequence[2], with the $^{13}$C carrier frequency set at 40 ppm and a duration of 1.44 ms, followed by $^{13}$C chemical shift evolution. During PC4, the $^{1}$H decoupling power was 70 kHz, and the $^{13}$C RF field strength was 5.5 kHz. The final $^{13}$C-$^{1}$H transfer step utilized an INEPT sequence[3] to specifically transfer polarization to the directly bonded protons, thereby enabling the assignment of the $^{1}$H chemical shifts attached to $^{13}$C. The MAS rate was 40 kHz. The $^{1}$H/$^{15}$N CP contact time was set to 1000 μs, with a $^{1}$H radio frequency (rf) field strength of approximately 70 kHz and a $^{15}$N rf field strength of approximately 30 kHz to fulfill the Hartmann-Hahn condition. The $^{15}$N/$^{1}$H CP field strengths were essentially identical to those of the $^{1}$H/$^{15}$N CP, but the $^{15}$N/$^{1}$H CP contact time was set to 0.2 ms to ensure single-bond polarization transfer. A linear ramp of 80% to 100% was applied on the $^{1}$H channel. All WALTZ-16 decoupling power levels were set to 10 kHz (0.25 * ω_MAS). XY decoupling power was set to 50 kHz. Typical 90° pulse lengths were 2.2 μs ($^{1}$H), 4.5 μs ($^{13}$C), and 5.5 μs ($^{15}$N). (B) 3D hCCH experiments using pulse sequence of 3D hCCH-INEPT-TOBSY[4] was employed as a reference to evaluate performance of that using 3D hCCH-INEPT-PC4. In this experiments, $^{1}$H-$^{13}$C polarization transfer was achieved via an INEPT sequence, followed by $^{13}$C chemical shift evolution. $^{13}$C-$^{13}$C transfer was accomplished using a TOBSY sequence[5,6], with the $^{13}$C carrier frequency set at 40 ppm and a duration of 6.0 ms, followed by $^{13}$C chemical shift evolution. The final $^{13}$C-$^{1}$H transfer step utilized an INEPT sequence to specifically transfer polarization to the directly bonded protons. (C) Comparison of the signal-to-noise ration of 3D hCCH spectra by using hCCH-CP-PC4 (red) and hCCH-INEPT-TOBSY (green). Only 1D of both spectra were acquired with 16 scans,the direct dimension acquisition time of 20 ms and no window function applied. The signal-to-noise ratio of the 1D hCCH-CP-PC4 spectrum is 8 times that of the 1D hCCH-INEPT-TOBSY spectrum[4]. Consequently, the hCCH-CP-PC4 sequence was used for the subsequent 3D hCCH experiments for assignments of methyl proton chemical shifts.

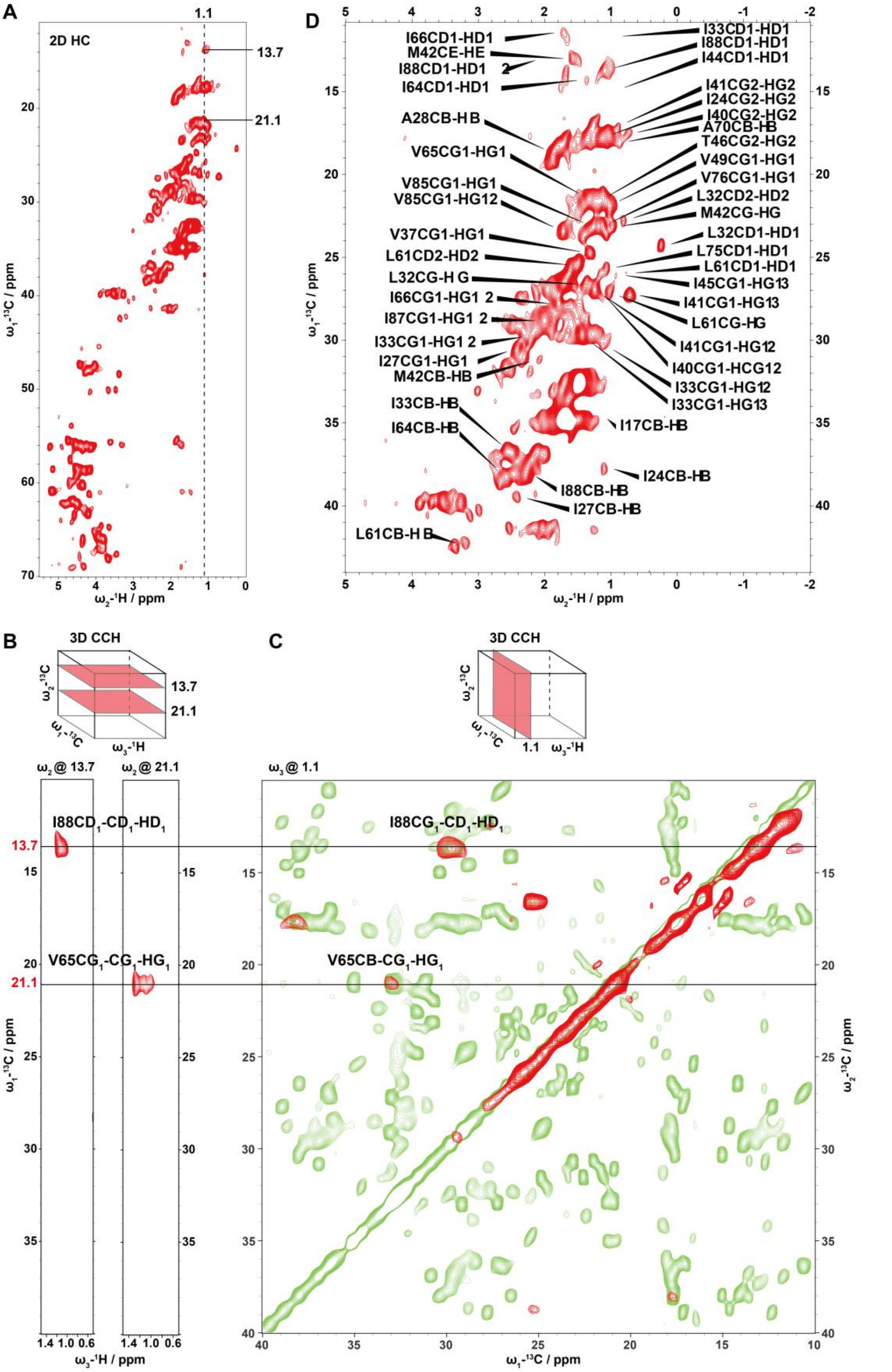


**Figure S3. Assignments of $^{1}$H chemical shifts of side-chain in MscL by using 3D hCCH spectra.** (A) 2D $^{1}$H-$^{13}$C CP-HSQC spectrum, with a zoomed-in region shown on the right. Assigned methyl proton chemical shifts are labeled on the spectrum. Detailed assignments are shown in **Table S2.** (B) The w1($^{13}$C)-w3($^{1}$H) plan extracted from the 3D hCCH spectrum at w2($^{13}$C) chemical shifts of 13.7 ppm and 21.1 ppm, showing the $^{13}$C–$^{1}$H assignments for residues I88 and V65. (C) The w1($^{13}$C)-w2($^{13}$C) plan extracted from the 3D hCCH spectrum at a w3($^{1}$H) chemical shift of 1.1 ppm, showing the $^{13}$C–$^{13}$C assignments for residues I88 and V65.

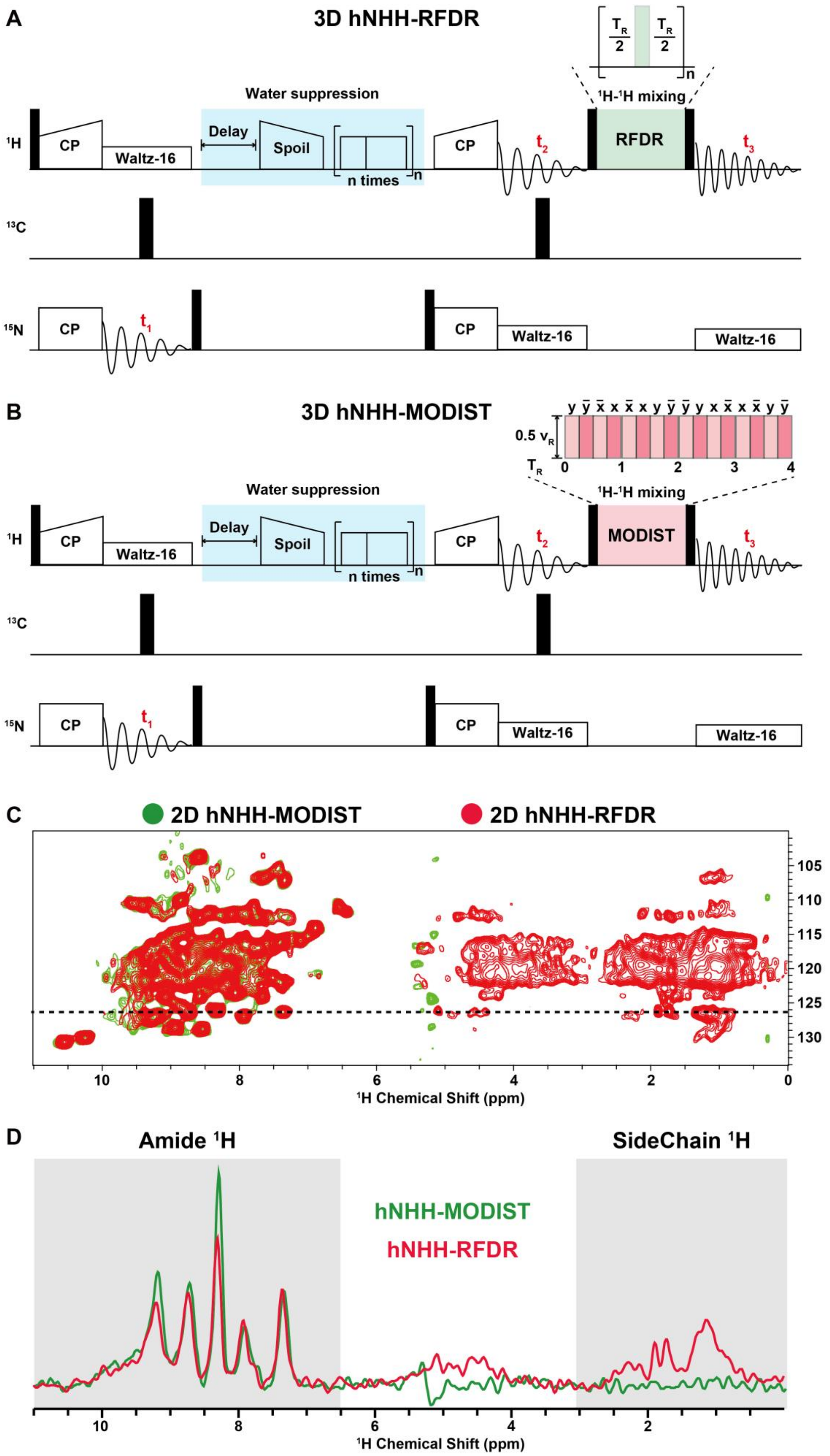


**Figure S4. 3D hNHH experiments for collecting distance constraints.** (A) Pulse sequence of 3D hNHH-MODIST was used in 3D hNHH experiments. In these 3D hNHH experiments, initial $^1H$-$^{15}N$ polarization transfer was achieved via cross-polarization (CP), followed by $^{15}N$ chemical shift evolution. The water suppression T-delay was set to 44 ms. The duration of the trapezoidal water suppression spoiler pulse was set to 1.2 ms, with its power level identical to the first $^1H$ CP pulse. The power level for the water suppression pulse train was set to 11 kHz, with pulse durations of 20 ms and 34 ms, respectively. Subsequently, $^{15}N$-$^1H$ polarization transfer was achieved via CP, followed by $^1H$ chemical shift evolution. $^1H$-$^1H$ transfer was accomplished using the MODIST recoupling module[7,8], followed by $^1H$ chemical shift evolution. For MODIST, the $^1H$ RF field strength was set to 20 kHz (0.5ωr), with a pulse width of 6.25us and recoupling times of 6 ms and 12 ms. (B) Pulse sequence of 3D hNHH-RFDR was used in 3D hNHH experiments. In these experiments, initial $^1H$-$^{15}N$ polarization transfer was achieved via cross-polarization (CP), followed by $^{15}N$ chemical shift evolution. The water suppression T-delay was set to 44 ms. The duration of the trapezoidal water suppression spoiler pulse was set to 1.2 ms, with its power level identical to the first $^1H$ CP pulse. The power level for the water suppression pulse train was set to 11 kHz, with pulse durations of 20 ms and 34 ms, respectively. Subsequently, $^{15}N$-$^1H$ polarization transfer was achieved via CP, followed by $^1H$ chemical shift evolution. $^1H$-$^1H$ transfer was accomplished using the RFDR recoupling module[9], with a pulse width of 5 μs and recoupling times of 2 ms and 4 ms, followed by $^1H$ chemical shift evolution. (C) Comparison of the sensitivity in 2D hNHH-MODIST (green) and 2D hNHH-RFDR (red) spectra of MscL in cellular membranes. (D) 1D $^1H$ traces extracted at a $^{15}N$ chemical shift of 126.3 ppm from the 2D hNHH-MODIST (green) and 2D hNHH-RFDR (red) spectra. The amide proton region of the hNHH-MODIST spectrum shows a 50% higher signal-to-noise ratio and exhibits no signals in the sidechain proton region, whereas the hNHH-RFDR spectrum shows signals in the side-chain proton region.

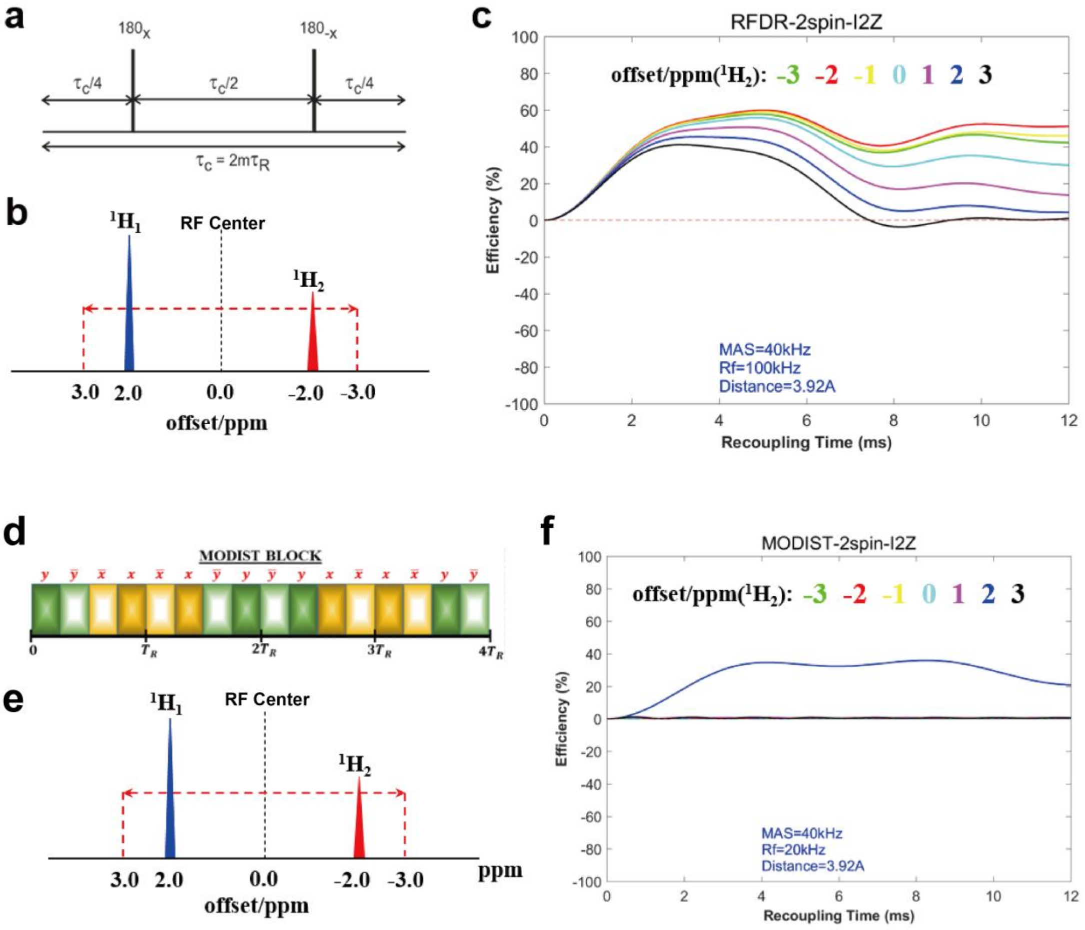


**Figure S5. Determination of optimal $^{1}$H-$^{1}$H mixing times for RFDR and MODIST recoupling modules through theoretical simulations.** (a-c) Simulations of the RFDR recoupling module using the Simpson program[10]. (a) Pulse sequence diagram of the RFDR recoupling module. (b) Simulation results of RFDR frequency selectivity, with $^{1}H_{1}$ as the target spin and $^{1}H_{2}$ as spins at different chemical shifts. It is observed that even when the chemical shift of $^{1}H_{2}$ varies (different colors represent different frequency offsets of $^{1}H_{2}$), the recoupling efficiency between $^{1}H_{1}$ and $^{1}H_{2}$ remains significant. (c) Theoretical simulations indicate that 2–4 ms is the optimal time window for RFDR recoupling. When the mixing time exceeds 4 ms, the intensity of $^{1}$H–$^{1}$H correlation signals shows significant attenuation. This attenuation is attributed to polarization leakage from backbone to sidechain protons, resulting from the combined effects of the broadband characteristics of RFDR and the presence of aliphatic protons in the [25% $^{1}$H,$^{13}$C,$^{15}$N]-MscL membrane sample. (d-f) Simulations of the MODIST recoupling module using the Simpson program. (d) Pulse sequence diagram of the MODIST recoupling module. (e) Simulation results of MODIST frequency selectivity, with $^1H_1$ as the target spin and $^1H_2$ as spins at different chemical shifts. The recoupling efficiency between $^1H_1$ and $^1H_2$ is found to vary significantly with the chemical shift of $^1H_2$ (different colors represent different frequency offsets of $^1H_2$). As shown in (e), high transfer efficiency from $^1H_1$ to $^1H_2$ is achieved only when $^1H_1$ and $^1H_2$ have identical chemical shifts. No magnetization transfer occurs between $^1H_1$ and $^1H_2$ spins at other chemical shifts, demonstrating the narrowband recoupling characteristic of MODIST. (f) Theoretical simulations show that MODIST requires mixing times longer than 4 ms to achieve optimal efficiency.

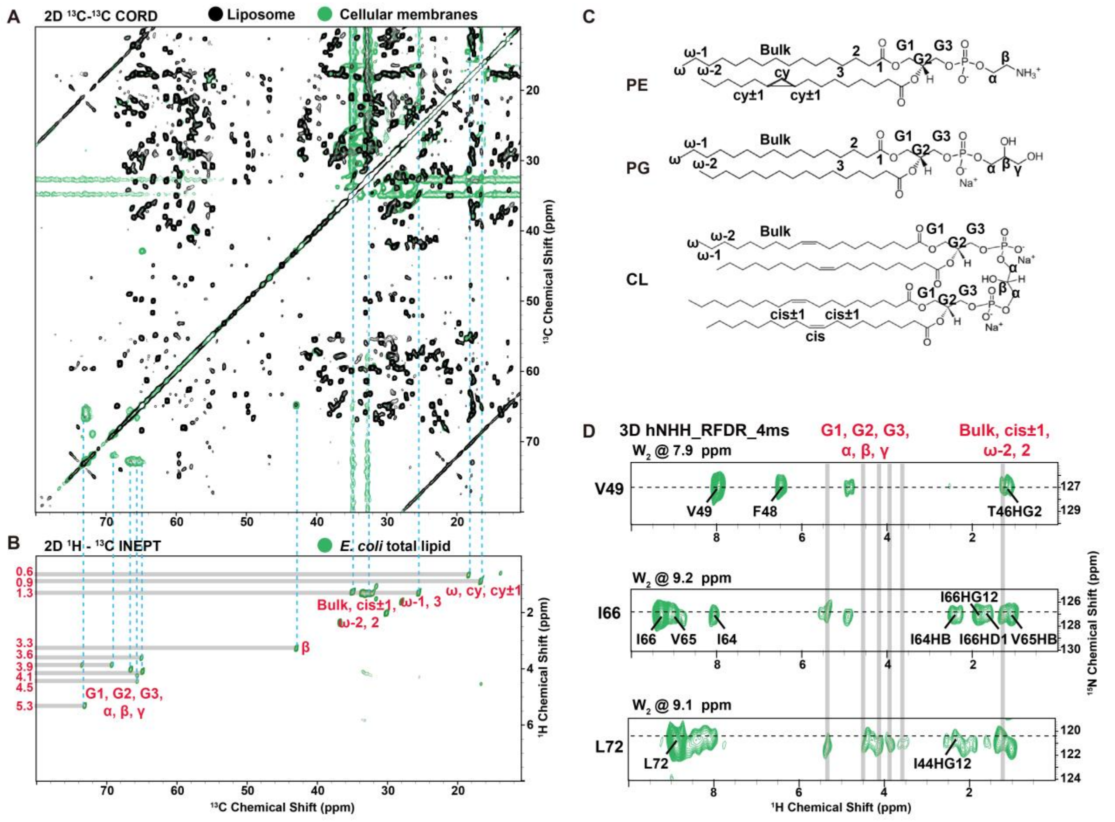


**Figure S6. Assignments of $^1$H chemical shifts of lipid signals in dipolar-based ssNMR spectra of MscL cellular membrane samples.** (A) Overlay of dipolar-based 2D $^{13}$C-$^{13}$C correlation spectra of *E. coli* lipids (black) and the MscL cellular membrane sample (green). (B) J-coupling-based 2D $^1$H-$^{13}$C spectrum (2D $^1$H-$^{13}$C INEPT) of the MscL cellular membrane sample. The $^{13}$C dimension is aligned with the dipolar-based 2D $^{13}$C-$^{13}$C spectrum to assign the $^1$H dimension of the 2D $^1$H-$^{13}$C INEPT spectrum. The assigned lipid proton chemical shifts are annotated in the spectrum. The corresponding lipid chemical shift assignments are summarized in **Table S3**. (C) Chemical structures and nomenclature of the major lipid types in *E. coli* lipids: phosphatidylethanolamine (PE), phosphatidylglycerol (PG), and cardiolipin (CDL). The phospholipid composition of E. coli is dominated by PE (69%), PG (19%), CDL (6.5%), and other minor species[11]. (D) Exclusion of the assigned lipid chemical shifts identified in (A) during the assignment of distance restraint peaks in the 3D hNHH spectrum.

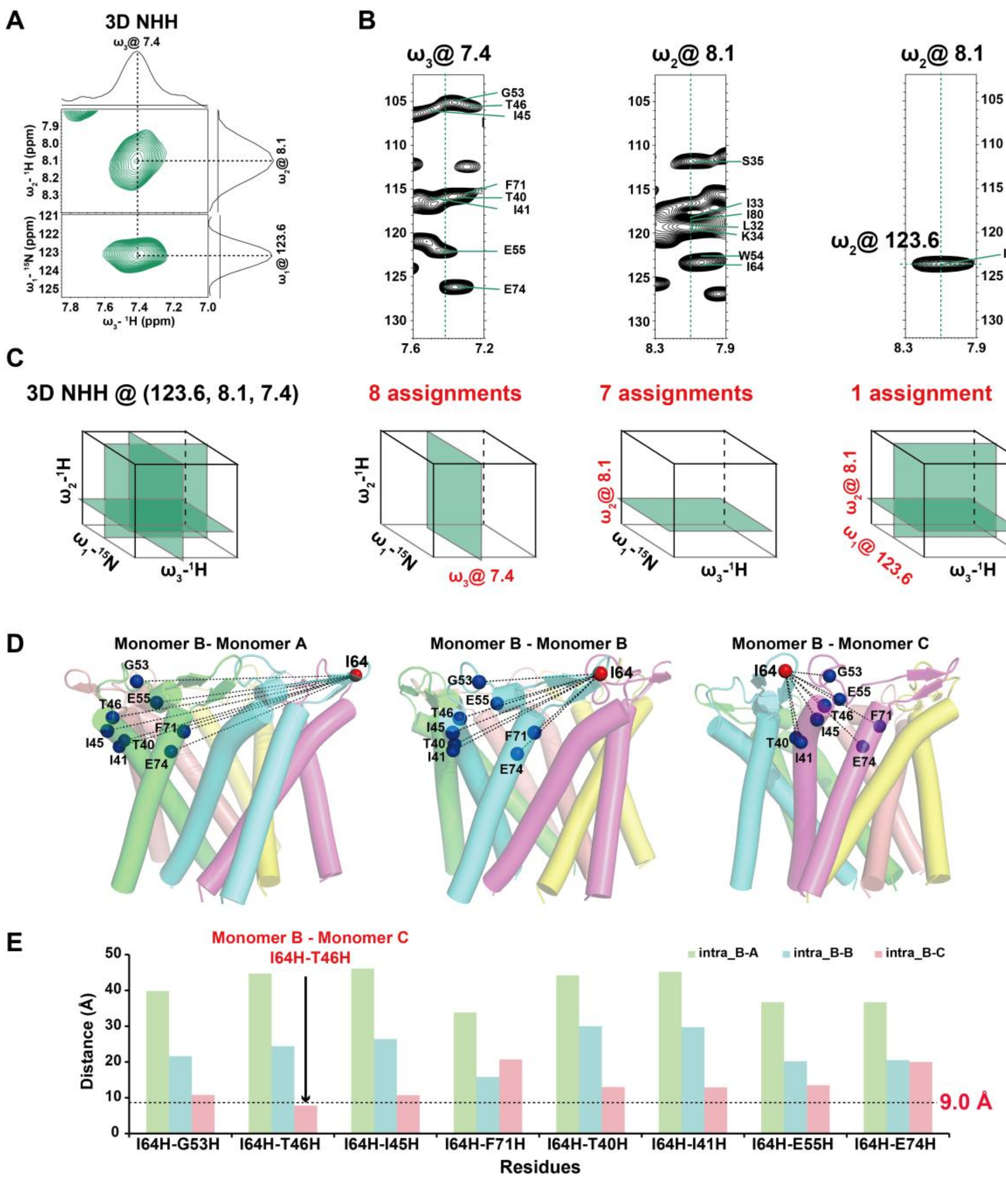


**Figure S7. Resolving ambiguity in assignment of spectral signals corresponding to distance restraints.** (A) Using the cross-peak at coordinates (123.6, 8.1, 7.4) ppm in the 3D hNHH spectrum to illustrate the ambiguity in assigning chemical shifts across different dimensions (w1, w2, w3). In the 2D NH spectrum, the w2 chemical shift at 8.1 ppm has seven possible assignments: S35, I33, I80, L32, K34, W54, and I64. The w3 chemical shift at 7.4 ppm has eight possible assignments: G53, T46, I45, F71, T40, I41, E55, and E74. Consequently, the cross-peak at (8.1, 7.4) ppm has 7 × 8 = 56 potential assignments. However, since w1 and w2 correspond to the amide nitrogen and proton within the same residue, respectively, the w1 chemical shift at 123.6 ppm unambiguously assigns the w2 signal at 8.1 ppm to I64, thereby reducing the assignment ambiguity by a factor of 7. (B) Chemical shifts (w1, w2, w3) from different dimensions of the 3D hNHH spectrum are displayed as solid green lines in the 2D NH spectrum. (C) The number of assignment possibilities shown in different 2D slices of the 3D NHH spectrum. (D) Validation of distances for four candidate assignment sites at the monomer B–monomer A interface, within monomer B, and at the monomer B–monomer C interface, using the pentameric structure derived from CS-Rosetta. (E) Distance distribution analysis: Only the intermolecular I64H–T46H pair satisfies the 9.0 Å distance threshold.

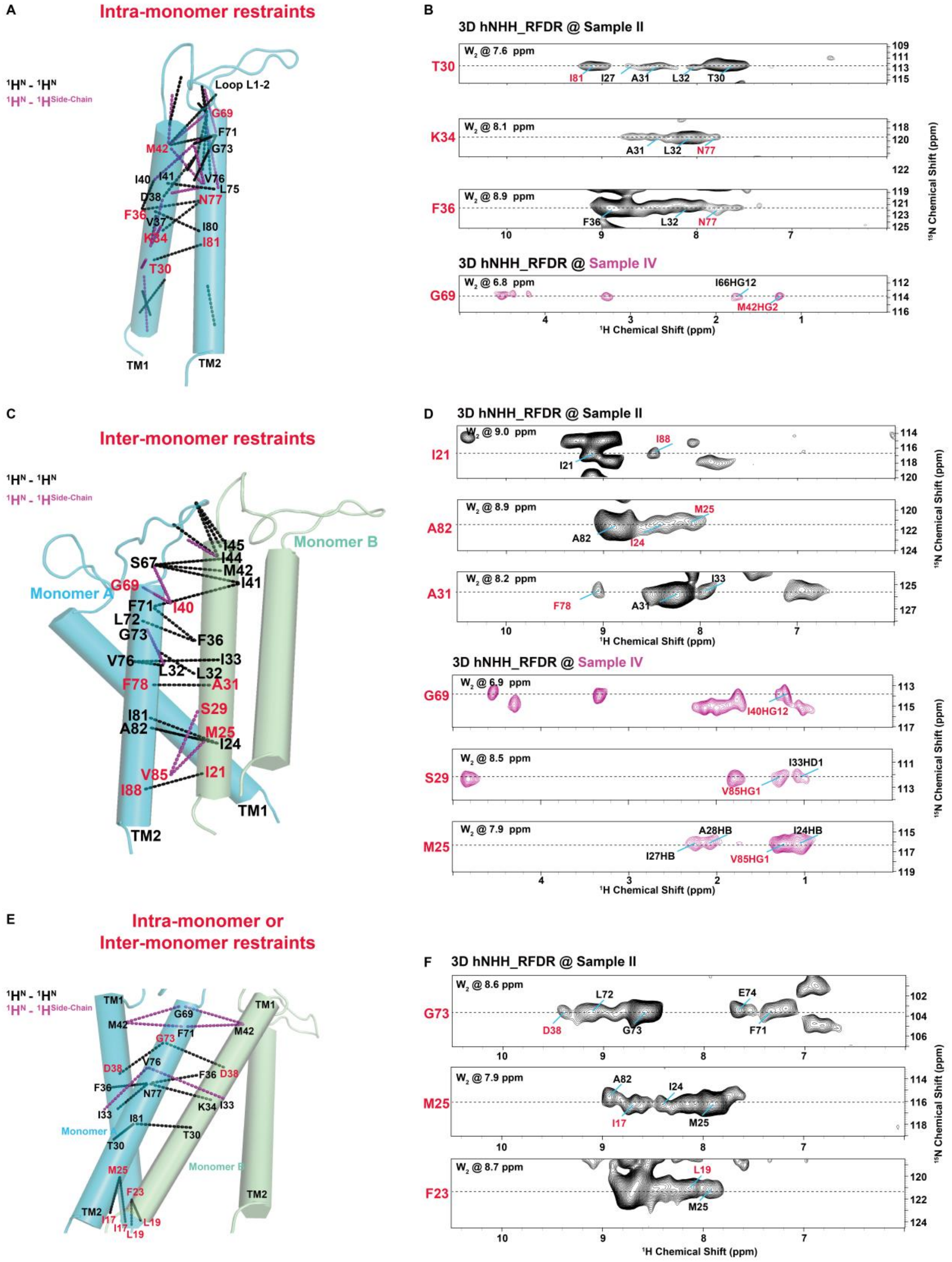


**Figure S8. All inter-helical long-range distance restraints extracted from ssNMR spectra.** (A) Mapping of all intra-subunit inter-helical distance restraints onto the MscL pentameric structure. Black lines represent $^1H^N$–$^1H^N$ restraints; purple lines represent $^1HN$–$^1H^{SC}$ restraints. (B) Key intra-subunit restraints shown in four 2D NH slices extracted from the 3D hNHH spectrum. (C) Mapping of all inter-subunit inter-helical distance restraints onto the MscL pentameric structure. Relevant residues are highlighted in red. Black spectra were acquired from Sample II; purple spectra were acquired from Sample IV. (D) Annotation of relevant residues on the MscL pentameric structure, with the residues of interest colored in red. (E) Mapping of all ambiguous inter-subunit inter-helical long-range distance restraints onto the MscL pentameric structure. Black lines indicate $^{1}$HN–$^{1}$HN restraints; purple lines indicate $^{1}H^{N}$–$^{1}H^{SC}$ restraints. (F) Three types of inter-subunit restraints displayed in three 2D NH slices extracted from the 3D hNHH spectrum. Relevant residues are marked in red. Black spectra were obtained using Sample II.

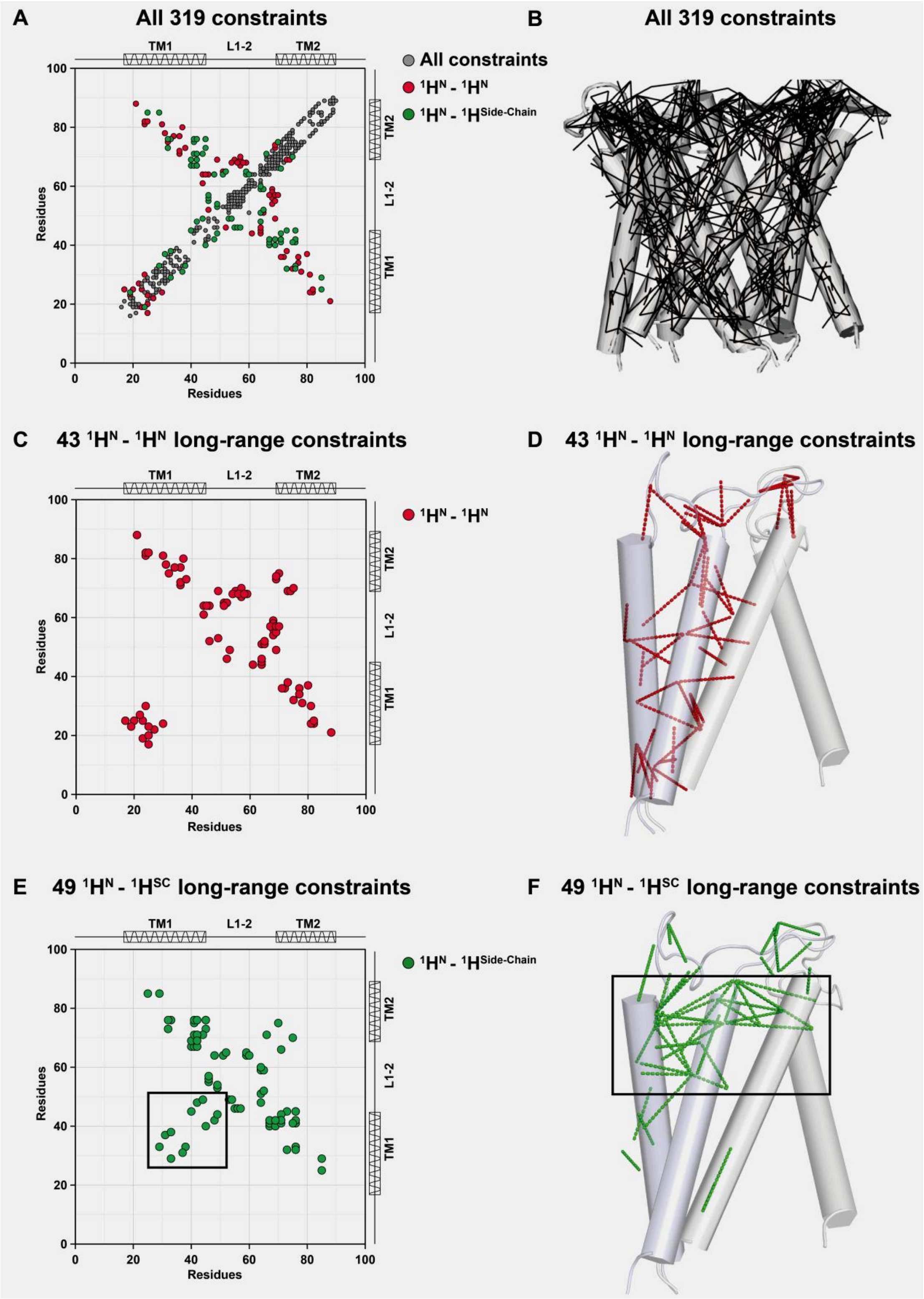


**Figure S9. Distance restraints extracted from ssNMR spectra of MscL in native cellular membranes and their structural implications, highlighting the critical role of side-chain restraints.** (A) Contact map displaying all 319 inter- and intra-subunit $^1H$–$^1H$ distance restraints within the MscL pentameric structure.

Cross-peaks are color-coded according to assignment type: $^1H^N$–$^1H^N$ in red, $^1H^N$–$^1H^{SC}$ in green. Secondary structure elements are indicated above the contact map. (B) All distance restraints mapped as black lines onto the MscL pentameric structure. (C) Contact map showing all 43 inter- and intra-subunit $^1$HN–$^1$HN distance restraints within the MscL pentameric structure. (D) All distance restraints mapped as red lines onto the MscL pentameric structure. (E) Contact map showing all 49 inter- and intra-subunit $^1H^N$–$^1H^{SC}$ distance restraints within the MscL pentameric structure. The red box highlights the additional restraints present in this set compared to the $^1H^N$–$^1H^N$ distance restraints. (F) All distance restraints mapped as green lines onto the MscL pentameric structure. The red box indicates the additional restraints relative to the $^1H^N$–$^1H^N$ distance restraints. These additional long-range $^1H^N$-$^1H^{SC}$ restraints are predominantly distributed between helices, thereby critically defining the helical packing geometry.

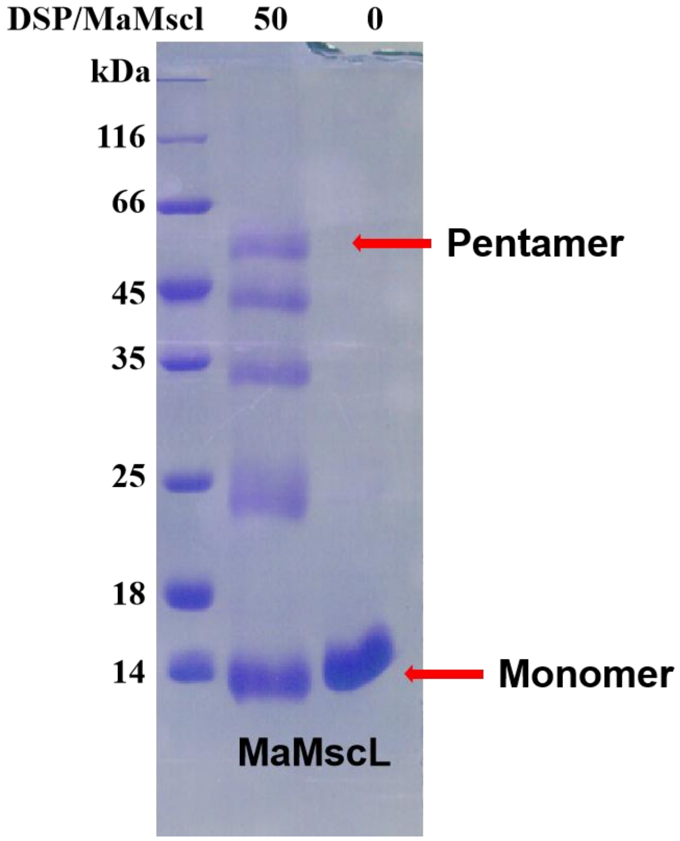


**Figure S10. Pentameric state of MscL confirmed by cross-linking experiments.** Direct determination of the oligomeric state of MscL in native membranes remains technically challenging due to steric hindrance from native lipids and limited accessibility of lysine residues within intact membrane microdomains[12]. Therefore, we solubilized cellular membranes and purified MscL into the detergent NSL to determine its aggregation state. In the cross-linking experiments, cross-linking agent of DSP was used with a molar ratio of DSP to MscL of 50. T The control on the right was not treated with DSP. After adding DSP to the MscL protein solution, the reaction was carried out at 37°C for 30 minutes and terminated by adding Tris solution. The results were analyzed by SDS-PAGE. The results showed that as DSP was added, the monomer content decreased while the pentamer content increased, indicating that MscL is a pentamer in detergents.

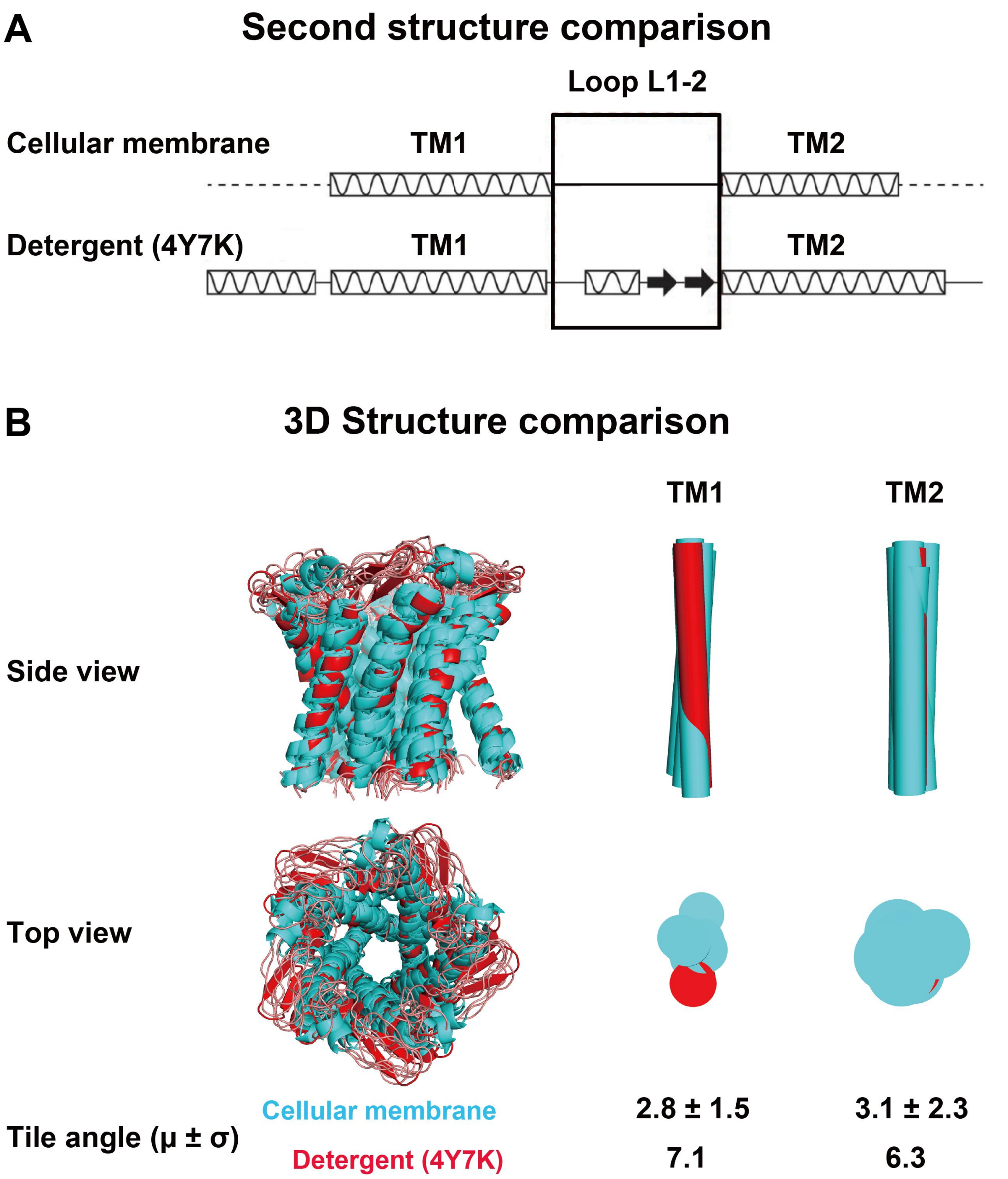


**Figure S11. Comparison of the structure determined by X-ray crystallography in detergent with the structure determined by *in situ* solid-state NMR in the membrane environment.** (A) Comparison of secondary structures. The secondary structure in the membrane environment was predicted from secondary chemical shifts[13]. The secondary structure in detergent was calculated from the crystal structure (PDB ID: 4Y7K)[14] using the DSSP program[15]. The results reveal subtle differences in the L1–2 loop region. (B) Comparison of 3D structures. The left panel shows side and top views of the crystal structure of the MscL pentamer determined in detergent (PDB

ID: 4Y7K) overlaid with the ensemble of 10 lowest-energy solid-state NMR structures determined in the native cellular membrane. The right panel illustrates differences in the tilt angles of the two transmembrane helices of MscL. The tilt angles were measured using PyMOL software relative to the average structure of the 10 lowest-energy MaMscL monomer structures in the native membrane, and are presented as mean ± standard deviation. Notably, the TM1 helix of MaMscL exhibits a significant difference in tilt angle (>2σ) between the native membrane and detergent environments.

**Table S1. Summary of solid-state NMR experiments and the used MscL samples.**

| ID | Samples | Experiment | $^1H$-$^1H$ mixing time | Indirect evolution | Scans per point | Experimental time |
|---|---|---|---|---|---|---|
| 1 | Sample II: *MscL* Cellular membrane Label: U- 25%$^1H$, $^{13}C$,$^{15}N$ Back exchange: 25%$H_2O$, 75%$D_2O$ | 2D $^1H$–$^{15}N$ CP-HSQC | - | 9.0 ms ($^{15}N$) | 64 | 2.4 h |
| | | 2D $^1H$–$^{13}C$ CP-HSQC | - | 7.0 ms ($^{13}C$) | 64 | 9.1 h |
| | | 3D hCCH_PC4 | - | 4.0 ms ($^{13}C$), 4.0 ms ($^{13}C$) | 16 | 103.4 h |
| | | 3D hNHH_RFDR_2ms | 2 ms | 8.0 ms ($^{15}N$), 5.0 ms ($^1H$) | 32 | 71.1 h |
| | | 3D hNHH_RFDR_4ms | 4 ms | 8.0 ms ($^{15}N$), 5.0 ms ($^1H$) | 48 | 133.3 h |
| | | 3D hNHH_MODIST_6ms | 6 ms | 8.0 ms (N), 5.0 ms ($^1H$) | 48 | 106.7 h |
| | | 3D hNHH_MODIST_12ms | 12 ms | 8.0 ms ($^{15}N$), 4.0 ms ($^1H$) | 64 | 113.8 h |
| 2 | Sample IV: *MscL* Cellular membrane Label: U- 10%$^1H$, $^{13}C$,$^{15}N$ Back exchange: 25%$H_2O$, 75%$D_2O$ | 2D $^1H$–$^{15}N$ CP-HSQC | - | 10.5 ms ($^{15}N$) | 64 | 2.1 h |
| | | 2D $^1H$–$^{13}C$ CP-HSQC | - | 7.0 ms ($^{13}C$) | 80 | 7.1 h |
| | | 3D hCCH_PC4 | - | 5.5 ms ($^{13}C$), 5.5 ms ($^{13}C$) | 32 | 142.2 h |
| | | 3D hNHH_RFDR_4ms | 4 ms | 9.7 ms ($^{15}N$), 5.0 ms (H) | 64 | 179.9 h |
| 3 | Sample I: *MscL* Cellular membrane Label: U- $^2H$, $^{13}C$,$^{15}N$ Back exchange: 100%$H_2O$ | 2D $^1H$–$^{15}N$ CP-HSQC | - | 9.0 ms ($^{15}N$) | 256 | 9.6 h |
| 4 | Sample III: *MscL* Cellular membrane Label: U- 10%$^1H$, $^{13}C$,$^{15}N$ Back exchange: 10%$H_2O$, 90%$D_2O$ | 2D $^1H$–$^{15}N$ CP-HSQC | - | 10.0 ms ($^{15}N$) | 128 | 6.4 h |
| | | 2D $^1H$–$^{15}N$ CP-HSQC_SWH | - | 9.0 ms ($^{15}N$) (SW_20kHz) | 96 | 14.4 h |
| 5 | *MscL* Cellular membrane Label: U- 5%$^1H$, $^{15}N$ Back exchange: 5%$H_2O$, 95%$D_2O$ | 2D $^1H$–$^{15}N$ CP-HSQC | - | 9.0 ms ($^{15}N$) (SW_20kHz) | 256 | 36.4 h |
| 6 | *MscL* E.coli lipids Label: U- 10%$^1H$, $^{13}C$,$^{15}N$ Back exchange: 10%$H_2O$, 90%$D_2O$ | 2D $^1H$–$^{15}N$ CP-HSQC | - | 9.0 ms ($^{15}N$) | 256 | 9.6 h |
| | | 2D $^1H$–$^{15}N$ CP-HSQC | - | 9.0 ms ($^{15}N$) (SW_20kHz) | 96 | 14.4 h |

**Table S2. Summary of side-chain proton chemical shift assignments of MscL in cellular membranes.**

| Residues | HB | HD1 | HD1 | HD2 | HE | HG | HG1 | HG1 | HG13 | HG2 |
|---|---|---|---|---|---|---|---|---|---|---|
| I17 | - | - | - | - | - | - | - | - | - | 1.076 |
| I24 | 1.058 | - | - | - | - | - | - | - | - | 1.105 |
| I27 | 2.401 | - | - | - | - | - | 2.45 | - | - | - |
| A28 | 1.869 | - | - | - | - | - | - | - | - | - |
| L32 | - | 0.242 | - | 0.79 | - | 1.44 | - | - | - | - |
| I33 | 2.361 | 0.968 | - | - | - | - | - | 2.184 | 1.463 | - |
| V37 | - | - | - | - | - | - | 1.36 | - | - | - |
| I40 | - | - | - | - | - | - | - | 1.242 | - | 0.730 |
| I41 | - | 0.705 | - | - | - | - | - | 1.245 | 0.670 | 1.041 |
| M42 | 2.047 | - | - | - | 1.66 | - | - | - | - | 1.106 |
| I44 | - | 1.059 | - | - | - | - | - | 2.518 | - | - |
| I45 | - | - | - | - | - | - | - | 1.151 | 0.967 | - |
| T46 | - | - | - | - | - | - | - | - | - | 1.183 |
| V49 | - | - | - | - | - | - | 1.09 | - | - | 1.313 |
| L61 | 3.24 | 0.781 | - | 1.59 | - | 1.10 | - | - | - | - |
| I64 | 2.47 | 1.356 | - | - | - | - | - | - | - | - |
| V65 | 1.115 | - | - | - | - | - | 1.39 | - | - | 1.182 |
| I66 | - | 1.514 | - | - | - | - | - | 1.826 | - | 1.330 |
| A70 | 0.830 | - | - | - | - | - | - | - | - | - |
| L72 | - | - | - | - | - | 1.28 | - | - | - | - |
| L75 | - | 1.189 | - | 1.78 | - | - | - | - | - | - |
| V76 | - | - | - | - | - | - | 1.15 | - | - | - |
| I81 | 1.918 | - | - | - | - | - | - | - | - | - |
| V85 | - | - | - | - | - | - | 1.24 | - | - | - |
| I88 | 2.263 | 1.406 | 1.924 | - | - | - | - | - | 1.115 | - |

**Table S3. Summary of proton chemical shifts of lipid signals present in dipolar-based spectra in MscL cellular membrane samples.**

| Number | Chemical Shift of Peak | | Assignments |
|---|---|---|---|
| | $^{1}H$ / (ppm) | $^{13}C$ / (ppm) | |
| 1 | 0.6 | 14.39 | ω, cy, cy±1 |
| 2 | 0.9 | 17.21 | |
| 3 | 1.3 | 32.13 | Bulk, *cis*±1, ω-1, 3, ω-2, 2 |
| 4 | 3.3 | 43.49 | β |
| 5 | 3.6 | 65.55 | α, β, γ, G1, G2, G3 |
| 6 | 3.9 | 69.82 | |
| 7 | 4.1 | 65.41 | |
| 8 | 4.5 | 66.17 | |
| 9 | 5.3 | 73.66 | |

**Table S4. List of all long-range distance constraints of MscL in cellular membranes.**

| Categories | | Number | Degeneracy | Distance restraints |
|---|---|---|---|---|
| Intra-Monomer | $^1H^N$-$^1H^N$ | 1 | 1 | T30N-H-I81H |
| | | 2 | 1 | T46N-H-G52H |
| | | 3 | 1 | A57N-H-G69H |
| | | 4 | 1 | W68N-H-W54H |
| | | 5 | 1 | A20N-H-M25H |
| | | 6 | 1 | V49N-H-G53H |
| | | 7 | 1 | A57N-H-S67H |
| | | 8 | 1 | I27N-H-A22H |
| | | 9 | 1 | W68N-H-T56H |
| | | 10 | 1 | W68N-H-V59H |
| | | 11 | 1 | V49N-H-G69H |
| | | 12 | 1 | E55N-H-G69H |
| | | 13 | 1 | A57N-H-G69H |
| | | 14 | 1 | A57N-H-A70H |
| | | 15 | 1 | W68N-H-T58H |
| | | 16 | 1 | A70N-H-L75H |
| | | 17 | 1 | G73N-H-G69H |
| | | 18 | 1 | E74N-H-G69H |
| | | 19 | 1 | K34N-H-N77H |
| | $^1H^N$-$^1H^{SC}$ | 20 | 1 | L19N-H-I24HB |
| | | 21 | 1 | I40N-H-I45HG12 |
| | | 22 | 1 | D38N-H-I33HB |
| | | 23 | 1 | F48N-H-M42HB |
| | | 24 | 1 | G53N-H-V49HG1 |
| | | 25 | 1 | V49N-H-I44HB |
| | | 26 | 1 | E55N-H-T46HG2 |
| | | 27 | 1 | T56N-H-T46HG2 |
| | | 28 | 1 | A57N-H-T46HG2 |
| | | 29 | 1 | V59N-H-I64HG12 |
| | | 30 | 1 | V59N-H-V65HG1 |
| | | 31 | 1 | E60N-H-I64HG12 |
| | | 32 | 1 | A70N-H-L75HD1 |
| | | 33 | 1 | F71N-H-I66HG12 |
| | | 34 | 1 | V76N-H-I41HG2 |
| | | 35 | 1 | V76N-H-M42HB |
| | | 36 | 1 | W54N-H-V49HG1 |
| | | 37 | 1 | A31N-H-V37HG1 |
| | | 38 | 1 | S29N-H-I33HD1 |
| | | 39 | 1 | V76N-H-I45HG1 |
| | | 40 | 1 | I41N-H-L75HD11 |
| | | 41 | 1 | S67N-H-I40HG12 |
| | | 42 | 1 | S67N-H-I41HG12 |
| | | 43 | 1 | S67N-H-M42HE1 |
| | | 44 | 1 | S67N-H-M42HB1 |
| | | 45 | 1 | G73N-H-I45HG23 |
| | | 46 | 1 | V76N-H-I41HG21 |

| Categories | | Number | Degeneracy | Distance restraints |
|---|---|---|---|---|
| | | 47 | 1 | V76N-H-M42HB1 |
| | | 48 | 1 | V76N-H-I45HG11 |
| | | 49 | 1 | V76N-H-I45HG23 |
| Inter-Monomer | $^1H^N$-$^1H^N$ | 1 | 1 | I21N-H-I88H |
| | | 2 | 1 | T46N-H-I64H |
| | | 3 | 1 | I64N-H-I44H |
| | | 4 | 1 | I64N-H-I45H |
| | | 5 | 1 | V65N-H-G52H |
| | | 6 | 1 | L61N-H-I44H |
| | | 7 | 1 | L32N-H-L75H |
| | | 8 | 1 | F36N-H-F71H |
| | | 9 | 1 | G51N-H-V65H |
| | | 10 | 1 | G52N-H-V65H |
| | | 11 | 1 | I64N-H-G51H |
| | | 12 | 1 | I24N-H-T30H |
| | | 13 | 1 | I24N-H-I81H |
| | | 14 | 1 | M25N-H-A82H |
| | | 15 | 1 | L72N-H-F36H |
| | | 16 | 1 | I80N-H-V37H |
| | | 17 | 1 | A82N-H-I24H |
| | | 18 | 1 | A82N-H-M25H |
| | | 19 | 1 | I88N-H-I21H |
| | $^1H^N$-$^1H^{SC}$ | 20 | 1 | M25N-H-V85HG1 |
| | | 21 | 1 | S29N-H-V85HG1 |
| | | 22 | 1 | F48N-H-I64HB |
| | | 23 | 1 | G51N-H-I64HG12 |
| | | 24 | 1 | G52N-H-V65HB |
| | | 25 | 1 | S67N-H-I40HG12 |
| | | 26 | 1 | G69N-H-I40HG12 |
| | | 27 | 1 | S67N-H-I41HG12 |
| | | 28 | 1 | S67N-H-M42HB1 |
| | | 29 | 1 | S67N-H-M42HE |
| | | 30 | 1 | G69N-H-I40HG12 |
| | | 31 | 1 | F71N-H-I41HG12 |
| | | 32 | 1 | F71N-H-I44HG12 |
| Inter-Monomer or Intar-Monomer | $^1H^N$-$^1H^N$ | 1 | 2 | A31N-H-F78H |
| | | 2 | 2 | F36N-H-N77H |
| | | 3 | 2 | K34N-H-N77H |
| | | 4 | 2 | M25N-H-I17H |
| | | 5 | 2 | F23N-H-L19H |
| | $^1H^N$-$^1H^{SC}$ | 6 | 2 | V76N-H-I33HB |
| | | 7 | 2 | V76N-H-L32HG |
| | | 8 | 2 | G73N-H-D38H |
| | | 9 | 2 | G73N-H-L32HG |
| | | 10 | 2 | F71N-H-M42HB1 |
| | | 11 | 2 | G69N-H-M42HB |

**Table S5. Summary of distance constraints from different experiments.**

| **Experiment Type** | **2ms hNHH_ RFDR** | **4ms hNHH_ RFDR** | **6ms hNHH_ MODIST** | **12ms hNHH_ MODIST** | **4ms hNHH_ Rfdr_ 90%RAP** | **Sum** |
|---|---|---|---|---|---|---|
| **Intra-residue restraints** | 29 | 21 | 32 | 2 | 0 | 84 |
| **Sequential restraints (\|i – j\| = 1)** | 24 | 27 | 21 | 7 | 0 | 79 |
| **Medium range restraints (2 ≤ \|i – j\| ≤ 4)** | 9 | 15 | 31 | 9 | 0 | 64 |
| **Long range restraints (\|i – j\| ≥ 5)** | 12 | 6 | 5 | 4 | 22 | 49 |
| **Inter-monomer restraints** | 8 | 6 | 10 | 8 | 11 | 43 |
| **Sum** | 82 | 75 | 99 | 30 | 33 | 319 |

**Table S6. Statistics for the structure determination of MscL**

| | MscL |
|---|---|
| **NMR distance and dihedral constraints** | |
| **Distance constraints** | |
| **Total NOE** | 362 × 5 |
| **Intra-residue** | 84 × 5 |
| **Inter-residue** | 278 × 5 |
| **Sequential (\|i – j\| = 1)** | 79 × 5 |
| **Medium range (2 ≤ \|i – j\| ≤ 4)** | 64 × 5 |
| **Long range (\|i – j\| ≥ 5)** | 49 × 5 |
| **Intermolecular** | 43 × 5 |
| **Hydrogen bonds** | 43 × 5 |
| **Total dihedral-angle restraints** | |
| **ϕ** | 67 × 5 |
| **ψ** | 67 × 5 |
| **Structure statistics** | |
| **Violations (mean ± s.d.)** | |
| **Distance constraints (Å)** | 0.003 ± 0.002 |
| **Dihedral-angle constraints (°)** | 1.466 ± 0.182 |
| **Max dihedral-angle violation (°)** | 8.94 |
| **Max distance-constraint violation (Å)** | 0.98 |
| **Deviations from idealized geometry** | |
| **bond lengths (Å)** | 0.003 ± 0.001 |
| **bond angles (°)** | 0.457 ± 0.013 |
| **Impropers (°)** | 0.354 ± 0.015 |
| **Average pairwise r.m.s.d. (Å)[a]** | |
| **Dimer structures** | |
| **Heavy** | 2.8 ± 0.6 |
| **backbone** | 1.9 ± 0.4 |
| **Structural quality** | |
| **Ramachandran Plot Statistics [b]** | |
| **Residues in most favored region (%)** | 92.5 |
| **Residues in additional allowed region (%)** | 7.5 |
| **Residues in generously allowed region (%)** | 0.0 |
| **Residues in disallowed region (%)** | 0.0 |
| **Clashscore[c]** | 13 |
| **Molprobity score (Å)** | 3.91 |

[a]Pairwise r.m.s. deviation was calculated between 10 structures, for residues in α-helices, including residues 17-46 and 67-90.

[b]Evaluated with the program PROCHECK[16].

[c]Evaluated with the Molprobity program[17].

### References


(1) Baldus, M.; Petkova, A. T.; Herzfeld, J.; Griffin, R. G. Cross Polarization in the Tilted Frame: Assignment and Spectral Simplification in Heteronuclear Spin Systems. *Mol, Phys,* **1998**, *95* (6), 1197–1207.

(2) Xiao, H.; Zhao, W.; Zhang, Y.; Kang, H.; Zhang, Z.; Yang, J. Selective Correlations between Aliphatic 13C Nuclei in Protein Solid-State NMR. *J. Magn. Reson.* **2024**, *365*, 107730.

(3) Morris, G. A.; Freeman, R. Enhancement of Nuclear Magnetic Resonance Signals by Polarization Transfer. *J. Am. Chem. Soc.* **1979**, *101* (3), 760–762.

(4) Agarwal, V.; Reif, B. Residual Methyl Protonation in Perdeuterated Proteins for Multi-Dimensional Correlation Experiments in MAS Solid-State NMR Spectroscopy. *J. Magn. Reson.* **2008**, *194* (1), 16–24.

(5) Baldus, M.; Meier, B. H. Total Correlation Spectroscopy in the Solid State. The Use of Scalar Couplings to Determine the through-Bond Connectivity. *J. Magn. Reson., Ser. A* **1996**, *121* (1), 65–69.

(6) Baldus, M.; Iuliucci, R. J.; Meier, B. H. Probing Through-Bond Connectivities and through-Space Distances in Solids by Magic-Angle-Spinning Nuclear Magnetic Resonance. *J. Am. Chem. Soc.* **1997**, *119* (5), 1121–1124.

(7) Nimerovsky, E.; Stampolaki, M.; Varkey, A. C.; Becker, S.; Andreas, L. B. Analysis of the MODIST Sequence for Selective Proton–Proton Recoupling. *J. Phys. Chem. A* **2025**, *129* (1), 317–329.

(8) Nimerovsky, E.; Najbauer, E. E.; Movellan, K. T.; Xue, K.; Becker, S.; Andreas, L. B. Modest Offset Difference Internuclear Selective Transfer via Homonuclear Dipolar Coupling. *J. Phys. Chem. Lett.* **2022**, *13* (6), 1540–1546.

(9) Bennett, A. E.; Rienstra, C. M.; Griffiths, J. M.; Zhen, W.; Lansbury, Jr., Peter T.; Griffin, R. G. Homonuclear Radio Frequency-Driven Recoupling in Rotating Solids. *J. Chem. Phys.* **1998**, *108* (22), 9463–9479.

(10) Bak, M.; Rasmussen, J. T.; Nielsen, N. Chr. SIMPSON: A General Simulation Program for Solid-State NMR Spectroscopy. *Magn. Moments* **2011**, *213* (2), 366–400.

(11) Morein, S.; Andersson, A.-S.; Rilfors, L.; Lindblom, G. Wild-Type Escherichia Coli Cells Regulate the Membrane Lipid Composition in a “Window” between Gel and Non-Lamellar Structures (∗ ). *J. Biol. Chem.* **1996**, *271* (12), 6801–6809.

(12) Merezhko, M.; Pakarinen, E.; Uronen, R.-L.; Huttunen, H. J. Live-Cell Monitoring of Protein Localization to Membrane Rafts Using Protein-Fragment Complementation. *Biosci. Rep.* **2020**, *40* (1), BSR20191290.

(13) Wishart, D. S.; Sykes, B. D. The 13C Chemical-Shift Index: A Simple Method for the Identification of Protein Secondary Structure Using 13C Chemical-Shift Data. *J. Biomol. NMR* **1994**, *4* (2), 171–180.

(14) Li, J.; Guo, J.; Ou, X.; Zhang, M.; Li, Y.; Liu, Z. Mechanical Coupling of the Multiple Structural Elements of the Large-Conductance Mechanosensitive Channel during Expansion. *Proc. Natl. Acad. Sci.* **2015**, *112* (34), 10726–10731.

(15) Kabsch, W.; Sander, C. Dictionary of Protein Secondary Structure: Pattern Recognition of Hydrogen-Bonded and Geometrical Features. *Biopolymers* **1983**, *22* (12), 2577–2637.

(16) Laskowski, R. A.; MacArthur, M. W.; Moss, D. S.; Thornton, J. M. PROCHECK: A Program to Check the Stereochemical Quality of Protein Structures. *J. Appl, Crystallogr,* **1993**, *26* (2), 283–291.

(17) Chen, V. B.; Arendall, W. B.; Headd, J. J.; Keedy, D. A.; Immormino, R. M.; Kapral, G. J.; Murray, L. W.; Richardson, J. S.; Richardson, D. C. MolProbity: All-Atom Structure Validation for Macromolecular Crystallography. *Acta, Crystallogr, D. Biol, Crystallogr,* **2010**, *66* (Pt 1), 12–21.
